# Gas diffusion kinetics drive embolism spread in angiosperm xylem: evidence from flow-centrifuge experiments and modelling

**DOI:** 10.1101/2023.04.19.537442

**Authors:** Luciano M. Silva, Luciano Pereira, Lucian Kaack, Xinyi Guan, Christophe L. Trabi, Steven Jansen

## Abstract

Understanding xylem embolism formation is challenging due to dynamic changes and multiphase interactions in conduits. If embolism spread involves gas movement in xylem, we hypothesise that it is affected by time. We measured hydraulic conductivity (K_h_) in flow-centrifuge experiments over one hour at a given pressure and temperature for stem samples of three angiosperm species. Temporal changes in K_h_ at 5, 22, and 35°C, and at various pressures were compared to modelled gas pressure changes in a recently embolised vessel in the centre of a centrifuge sample. Temporal changes in K_h_ at 22°C showed maximum relative increases between 6% and 40%, and maximum decreases between 41% and 61% at low and high centrifugal speed, respectively. Logarithmic changes in K_h_ were species-specific, and most pronounced during the first 15 minutes. Embolism formation started near the edges of centrifuge samples and gradually increased at the centre. Moreover, measured decreases in K_h_ strongly correlated with modelled increases in gas concentration in a recently embolised vessel. Although embolism is mostly pressure-driven, our experimental and modelled data indicate that time, conduit characteristics, and temperature are involved due to their role in gas diffusion. Gas diffusion, however, does not cover the entire process of embolism spread.

## Introduction

During drought stress, the pressure of xylem sap can become negative enough to allow a local phase change from liquid to water vapour (Tyree & Zimmermann, 2002). If this phase change takes place, the water vapour, which is gradually mixed with gas, fills the entire water-conducting conduit and blocks the water transport by a process known as embolism (Hölttä *et al*., 2002; Wang *et al*., 2015). There is growing evidence that embolism formation is induced in sap-filled conduits that are directly connected to an embolised vessel via a joint interconduit wall (Brodersen & McElrone, 2013; Knipfer *et al*., 2015; Choat *et al*., 2016; Skelton *et al*., 2017; Guan *et al*., 2021). Although embolism formation in xylem conduits has been given considerable research attention, we lack a mechanistic understanding of how exactly embolism is formed, and how it propagates between vessels (Jansen & Schenk, 2015; Levionnois *et al*., 2020; Lens *et al*., 2022; Lintunen *et al*., 2022). Bordered pits in interconduit walls are known to play a major role in transport of liquids and gasses across conduits (Kaack *et al*., 2019, 2021; Park *et al*., 2019; Zhang *et al*., 2020). Therefore, the ultrastructure of the pit membrane in bordered pit pairs of adjacent conduits is essential for fluid transport and embolism resistance.

It has also become clear that embolism resistance is a crucial component of plant-water relations, and is mainly associated with drought-induced plant mortality (Brodribb *et al*., 2010, 2021; Anderegg *et al*., 2012; Choat *et al*., 2012). Once embolism has reduced the number of functional conduits in xylem, the sap transport capacity is decreased until new functional conduits have been developed (Tyree & Sperry, 1989). Thus, considerable levels of embolism reduce the plant’s water supply, with consequences for photosynthesis. Under persistent, recurrent, or increasing periods of drought, embolism may ultimately contribute to desiccation, loss of organs, and partial or complete plant die-back (Tyree & Sperry, 1988; Anderegg *et al*., 2016).

The mechanism behind embolism formation has traditionally been described as “air-seeding”, and the Young-Laplace equation has been used to predict the pressure difference that a gas bubble would require to penetrate a pore in a pit membrane by mass flow and then induce embolism (Tyree & Zimmermann, 2002). However, the Young-Laplace equation is inappropriate when considering the complex gas-liquid interfaces inside a three-dimensional pit membrane, which shows multiple pore constrictions in each pore pathway (Kaack *et al*., 2021). Although embolism formation seems to be largely a pressure-driven process, with more negative pressure of xylem sap increasing the likelihood of embolism (Jansen & Schenk, 2015; Levionnois *et al*., 2020; Lens *et al*., 2022; Lintunen *et al*., 2022), there is evidence suggesting that embolism formation may not only be pressure-driven. Indeed, embolism resistance may also depend on the proximity to a gas source and local changes in gas pressure and concentration, which could lead to spreading of embolism at different xylem water potentials (Choat *et al*., 2010; Torres-Ruiz *et al*., 2015; Guan *et al*., 2021; Avila *et al*., 2022).

The level of xylem embolism can be indirectly quantified by measuring the reduction of the hydraulic conductivity (K_h_) when a xylem sample is submitted to a decreasing xylem water potential (Sperry, 1985). Based on this concept, the flow-centrifuge is one of various methods to construct vulnerability curves within a relatively short time frame of ca. 30 minutes to 1 hour (Cochard, 2002; Cochard *et al*., 2005, 2007). Although centrifuge methods have been used in many studies (Cochard *et al*., 2013), it remains unclear how exactly embolism inside a centrifuge sample is generated, and how vessel dimensions may affect embolism formation (Cai *et al*., 2010).

It is known that any hydraulic measurement is affected by temperature as it has an impact on conductance, modifying considerably the liquid viscosity, and also the cell membrane fluidity and permeability for symplastic transport (Cochard *et al*., 2000; Wang *et al*., 2014; Burlett *et al*., 2022a). Wang *et al*. (2014) reported that temperature affected K_h_ in a flow-centrifuge by 2.4% °C^-1^. Moreover, temperature has an influence on the gas solubility, and thus on the concentration of dissolved gas in xylem sap (Weiss, 1970). A potential link between the amount of gas dissolved in xylem sap and embolism formation has been suggested (Guan *et al*., 2021), but remains largely untested. If changes in the concentration of dissolved and undissolved gas would be associated with embolism formation, time may also affect the rate of embolism propagation because gas diffusion can be relatively slow from seconds to many hours (Sorz & Hietz, 2006; Yang *et al*., 2023). Thus, based on this assumption, it would be reasonable to expect that both time and temperature affect embolism formation in flow-centrifuge experiments. Interestingly, Wang *et al*. (2014) reported the challenge of predicting K_h_ using linear regression at the end of a vulnerability curve, suggesting that “new embolism may have been occurring during the process” (i.e. during spinning of a flow-centrifuge at a fixed speed). Also, K_h_ measurements over long time periods (hours) have been reported to show a constantly decreasing trend. This decrease in K_h_ has been explained by clogging or coating of intervessel pit membranes, wounding response, or another unknown phenomenon (Espino & Schenk, 2011; De Baerdemaeker *et al*., 2019). Therefore, it is important to test if any decline in K_h_ over time is caused by embolism propagation, related to a change in temperature, and/or caused by any other unknown factor.

There is rather contradictory information in literature about the distribution of embolised vessels along the axis of centrifuged samples. While some studies have shown that the highest level of embolism occurs in the centre (Cai *et al*., 2010; Sperry *et al*., 2012; Tobin *et al*., 2013), others have reported that embolism occurred mainly near the sample ends (Cochard *et al*., 2010; Martin-StPaul *et al*., 2014; Yin *et al*., 2019). Theoretically, the most negative pressure occurs at the centre of the rotor, with the pressure increasing along the axis towards the outer sides of the sample (Alder *et al*., 1997; Cai & Tyree, 2010; Cai *et al*., 2010). Therefore, the amount of embolism formed in the stem should be proportional to the pressure that is applied to the centre (Cochard *et al*., 2010; Yin *et al*., 2019). Other researchers observed that the highest embolism levels occur at the water injection side of samples in a flow-centrifuge (Martin-StPaul *et al*., 2014; López *et al*., 2018; Yin *et al*., 2019). An explanation that has been suggested is that tiny bubbles migrate with the flow, creating a gradient in loss of conductivity along the stem, with the greatest loss of conductivity at the basal and upstream part of the segment. There is also evidence suggesting that vessel dimensions play a role in embolism formation in the flow-centrifuge, with artificial embolism formation in conduits that are completely cut-open at both stem ends (Lamarque *et al*., 2018). Although we do not fully understand the relationship between vessel dimensions and vulnerability to embolism in plants (Lens *et al*., 2022), vessel dimensions play a role in the movement of gas between vessels (Salomón *et al*., 2021; Yang *et al*., 2023), and may therefore affect embolism location in a sample used for centrifuge-flow experiments.

This study aims to elucidate fundamental mechanisms behind embolism propagation in a flow-centrifuge. In particular, we hypothesize that if gas movement between conduits is relatively slow and related to embolism formation and propagation, there would be a considerable time effect on K_h_ measurements, and temporal changes in embolism location in flow-centrifuge experiments. By modelling how time, temperature and vessel dimensions affect gas movement and gas solubility, we expect that there is a correlation between the gas concentration in a recently embolised water vapour-filled vessel and the loss of K_h_ of xylem. Finally, we speculate that vessel dimensions affect the speed of gas movement in stem xylem, and therefore the rate and location of embolism in samples subject to spinning in a flow-centrifuge.

## Material and methods

### Plant material and site information

The temperate angiosperm tree species *Corylus avellana, Fagus sylvatica* and *Prunus avium* were selected. These are common trees near Ulm University (Germany, 48°25’20.3” N, 9°57’20.2” E).

### Measurements of hydraulic conductivity

Flow-centrifuge measurements were performed with a ChinaTron centrifuge (Model H2100R, Cence Company, Xiangyi, China) with manual control of the rotation speed and temperature. The ChinaTron flow centrifuge used had a thermostat temperature sensor near the rotor and a reversing valve. By actively heating or cooling the heat pump, the temperature was set with high precision to a particular value ± 0.1°C (Wang *et al*., 2014). Software designed by Yujie Wang recorded centrifuge parameters (temperature and rotational speed), calculated the water tension at the central axis of rotation, and recorded with a highspeed camera (scA640-120gm, Basler, Ahrensburg, Germany) changes in the positions of the menisci over time, which allowed us to estimate K_h_ based on Alder *et al*. (1997). The water tension at the central axis of rotation was used to estimate xylem water potential (Ψ) by subtracting an atmospheric pressure of 0.1 MPa.

The spinning duration of a centrifuge experiment was started as soon as a desired rotational speed was achieved. The acceleration and deceleration of the flow-centrifuge were set to 38.5 RPM s^-1^. Therefore, high spinning speeds required a somewhat longer acceleration time.

### Sample preparation

To avoid embolism formation by cutting branches with xylem sap under considerable negative pressure (Wheeler *et al*., 2013; Torres-Ruiz *et al*., 2015; Guan *et al*., 2021), long branch samples were first cut in air and immediately placed in a water-filled bucket. Then, the samples were transferred to the lab within 10 min, where the stem base was recut underwater. Finally, the samples were kept in water for more than 2 h for rehydration.

After sample rehydration, the samples were recut several times underwater to a final length of 27.4 cm (Torres-Ruiz *et al*., 2015). To fit the sample in the ChinaTron rotor (Cence Company, Xiangyi, China), we selected a plant segment with a diameter (excluding the bark) of 6 ±2 mm. Then, the stem segment ends were trimmed and the bark was removed with a sharp razor blade. The stem segments were mounted in the centrifuge rotor with their ends in cuvettes, following the natural orientation of xylem sap flow. Flow centrifuge measurements were performed with a solution of 10 mM KCl and 1 mM CaCl_2_ (Burlett *et al*., 2022b), which was injected with a peristaltic pump (Model PP 2201, VWR International bvba, Leuven, Belgium) into the cuvettes to measure K_h_.

*Experiment 1: Continuous vs. non-continuous flow in hydraulic measurements over a long spinning time* To test if the decline in K_h_ over time is caused by embolism propagation, and not by other factors, we assessed the effect of the solution injection on flow-centrifuge measurements of K_h_. To separate these effects, we spun samples under two different conditions: (1) continuous addition of the KCl and CaCl_2_ solution to the cuvettes to maintain an uninterrupted flow within the sample, and (2) no continuous addition of the solution (i.e. non-continuous flow) between K_h_ measurements. For continuous flow, the solution was injected in the cuvette continuously, keeping the menisci of the cuvette in movement, which means that a non-stop flow throughout the sample was applied. For non-continuous flow, the solution was injected only when we took a measurement. For *C. avellana*, K_h_ was assessed every 30 minutes for 8 hours at 22°C. The rotational speed of 3800 RPM was applied (Ψ = - 1.17 MPa), which caused less than 12% of embolism according to Guan *et al*. (2022b).

*Experiment 2: The effect of water potential, time and temperature on xylem embolism propagation*

In order to uncouple the effect of time, temperature, and Ψ on K_h_ measurements, we applied a given rotational speed and temperature to the ChinaTron, and took K_h_ measurements over the duration of one hour. Thus, one treatment consisted of a single sample that was spun in the centrifuge under a fixed speed and temperature. The experiment was repeated for three different temperatures (5 °, 22 °, and 35 °C), and several rotational speeds.

The temperature values selected were realistic temperatures that plants are exposed to in the field, and feasible with our ChinaTron system. For 5° and 35°C, the solution was manually injected into the cuvettes with a syringe. Before injection, the syringes were stored in a freezer and in a water bath, which were at 5°C and 35°C, respectively. Room temperature in the lab was 22°C. The temperature of the centrifuge solution was checked before injection with an electronic thermometer (16200, Bioblock Scientific, Illkirch, France).

For the 22°C condition, we applied different rotational speeds, namely eight for *F. sylvatica*, and nine for *C. avellana*, and *P. avium*. The speed values selected allowed us to link embolism propagation over time with different levels of initial embolism (Table S1). The two highest and lowest rotational speeds applied at 22°C were also used for measurements at 5° and 35°C (Table S1). Overall, the rotational speed, especially the highest ones, varied according to the species, and were selected based on vulnerability curves previously conducted on samples from the same species by Guan *et al*. (2022b).

K_h_ was measured several times over one hour. The time interval between consecutive measurements increased over time: every 20 seconds during the first 15 minutes of spinning, every minute during the next 15 minutes, and every 5 minutes during the next 15 minutes. A final measurement was taken when the sample had been spinning for 60 min. Therefore, this approach was optimized to capture the highest changes in K_h_ during the initial stages of the 1-hour long spinning experiments.

*Experiment 3: Location of embolised vessels in stem segments*

To assess the distribution of embolised vessels within 27.4 cm stem segments, we used an ultra-low flowmeter system, as proposed by Tyree et al. (2002) and adapted by Pereira & Mazzafera (2012). For each of the three species studied, we divided 12 samples into three groups (n = 4), and spun all samples in the ChinaTron at 22°C. To monitor potential changes in embolism over time, each group of samples was subjected to a different spinning time, which varied among the species: 1, 7.5 and 15 minutes of spinning for *C. avellana*; 1, 30 and 60 minutes for *F. sylvatica*, and 1, 15 and 30 minutes for *P. avium*. These arbitrary spinning times were selected based on trials (data not shown), which aimed to find the lowest spinning time differences that show changes in embolism level. The highest rotational speeds applied to each species in experiment 2 were used because the ultra-low flowmeter could not detect embolism in rotational speeds below these values. We prepared stem segments with cut-open vessels, which were required to remove bubbles in conduits by flushing the stem segments. The length of each stem segment was shorter than the mean vessel length for each species (Table S2 and S3).

After spinning, samples of *C. avellana, F. sylvatica* and *P. avium* were re-cut under water in 9 segments of 3.0 cm, 6 segments of 4.5cm, and 5 segments of 5.4 cm, respectively. After re-cutting, both ends of the segments were carefully trimmed with a sharp razor blade, connected in series to the hydraulic apparatus, and the initial conductance was measured (Pereira & Mazzafera, 2012). Then, embolism was removed by flushing each segment with a 6 ml syringe with a 10 mM KCl and 1 mM CaCl_2_ solution until bubbles were no longer observed at the distal stem end. Finally, the maximum conductivity of the segments was assessed. The Percentage Loss of Conductivity (PLC) was calculated for each of the segments using the hydraulic conductivity measurements taken before (K_0_’) and after (K_F_’) flushing the segments.

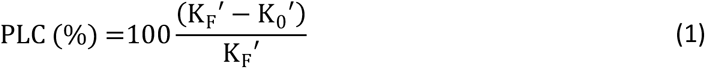

### Modelling of the gas pressure in a recently embolised vessel in flow-centrifuge experiments

Since we did not know how exactly embolism formation was trigged (Schenk *et al*., 2015; Kaack *et al*., 2021), this process could not be directly modelled. However, once a conduit has been embolised, it is known that this conduit is initially filled with 100% of water vapour, and gradually becomes filled with gas until atmospheric pressure has been achieved (Wang *et al*., 2015; Yang *et al*., 2023). As this process is associated with local gas transport and changes in pressure, we assumed that this step would be related to embolism resistance. Therefore, we developed a gas diffusion model to predict changes of the gas pressure in recently embolised conduits in a flow-centrifuge (Box S1). This model was based on a similar vessel geometry as developed for the Unit Pipe Pneumatic model (UPPn) of Yang *et al*. (2023), which estimated gas diffusion between intact conduits via interconduit pit membranes. However, our model estimated the gas dissolved in water-filled conduits and the gas concentration in embolised vessels. Considering the centrifuge force in the flow-centrifuge, the model estimated how anatomical parameters, temperature, time, and Ψ would affect the concentration of gas in a recently embolised vapour-filled vessel.

At a time zero (t=0), we simulated an embolised water vapour-filled vessel at the centre of a centrifuge sample, and modelled its relative changes in gas concentration over time (C_relative_). We assumed that the gas diffusion near the centre of the stem segment was through intervessel pit membranes only, mainly because gas diffusion across pit membranes is ca. 100 times faster than across cell walls (Yang *et al*., 2023). The model estimated the axial gas diffusion in sap from the stem ends through successive water-filled vessels towards the centre of the sample, and finally through a wet pit membrane into the embolised water-vapour-filled vessel. A list of all abbreviations, units and definitions used in this study can be found in Table S5 and S6. All equations and calculation steps were detailed in the Supporting Information (Methods S1).

### Statistics and data analysis

Data processing, simulations and statistical analyses were performed in RStudio 2022 (version 4.2.2, R Core Team, Boston, USA), OriginPro 2016 (version b9.3.226, OriginLab Corporation, Northampton, USA) and Excel 2021 (version 16.56, Microsoft, Redmond, USA) software.

To compare the species as well as the different conditions applied in Experiments 1 and 2, we used relative values of K_h_, defined here as the relative change in hydraulic conductivity over time (ΔK_h_, %), which was calculated as follows:

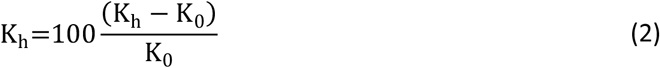

Where K_0_ was the hydraulic conductivity measured at time zero, and K_h_ was the current hydraulic conductivity value. Thus, ΔK_h_ described the temporal changes in K_h_ as relative values to its first value measured with a flow-centrifuge at a given spinning speed.

We used non-linear regressions to test how time, Ψ and their interaction affected ΔK_h_ for the centrifuge-flow measurements at 22°C in experiment 2. All the equations and calculation steps were described in detail in the Supporting Information (Methods S2).

Considering the two most negative Ψ values applied in experiment 2, we assessed the equation that described K_h_ as a function of time (K_h_[t]). For this, we fitted a rectangular hyperbola model based on Bliss & James (1966), and estimated the equation coefficients for each species and condition applied. All the equations and calculation steps were detailed in the Supporting Information (Methods S3). Based on the K_h_(t) equation, we estimated the minimum value of hydraulic conductivity (K_min_), and then calculated the relative hydraulic conductivity (K_relative_, in %) as follows:

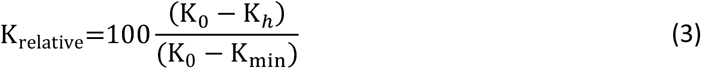

Assuming that the gas pressure in a recently embolised vessel was mechanistically related to K_h_, the linear relationship between C_relative_ and K_relative_ was assessed and used to evaluate the performance of our gas diffusion model. To quantify how much the model could predict xylem embolism propagation in a flow-centrifuge, we computed both the root mean squared error (RMSE) and Person correlation coefficient (R) of this relationship.

In experiment 3, a single stem segment was considered as one replication, and thus the spinning time and the location of the stem segment within the 27.4 cm long centrifuge sample were the sources of variation. The effects of these two factors and their interaction on PLC were tested by two-way analysis of variance (ANOVA). When significant effects were detected, the mean values were compared using Tukey’s HSD test at 95% confidence. We verified ANOVA’s assumption and, when necessary, transformed the data according to Dag & Ilk (2017).

## Results

### Experiment 1: Continuous vs. non-continuous flow in hydraulic measurements over a long spinning time

After 8 hours of spinning, changes in the percentage of the hydraulic conductivity (ΔK_h_) were 23.7% for the sample of *C. avellana* with continuous addition of solution (continuous flow), and 4.49% for the *C. avellana* sample with no solution addition (non-continuous flow) [Fig. S1]. While for continuous flow, a linear decrease of ΔK_h_ was noticed from the beginning (with ΔK_h_ decreasing 9.1% in the first 30 minutes), a minor decrease in ΔK_h_ of 0.21% was observed after 6 hours of spinning during non-continuous flow. Therefore, flow measurements were found to be stable for several hours of spinning under non-continuous flow condition.

### Experiment 2: The effect of water potential, time and temperature on xylem embolism propagation

At 22°C, K_h_ and ΔK_h_ changed logarithmically over time, decreasing for water potentials (Ψ) close to -4.00 MPa, but increasing when Ψ was close to zero (Fig. 1 and Fig.S2). From -0.25 to -0.83 MPa, ΔK_h_ increased over time for all three study species. For *C. avellana* and *F. sylvatica*, ΔK_h_ was stable and around zero when Ψ was -0.98 and -2.00 MPa, respectively. For *P. avium*, no speed applied resulted in stable ΔK_h_ values. Values of Ψ higher than -1.31, -2.11, and -2.87 MPa, resulted in a temporal increase of ΔK_h_ for *C. avellana, F. sylvatica*, and *P. avium* respectively.

**Fig. 1.**
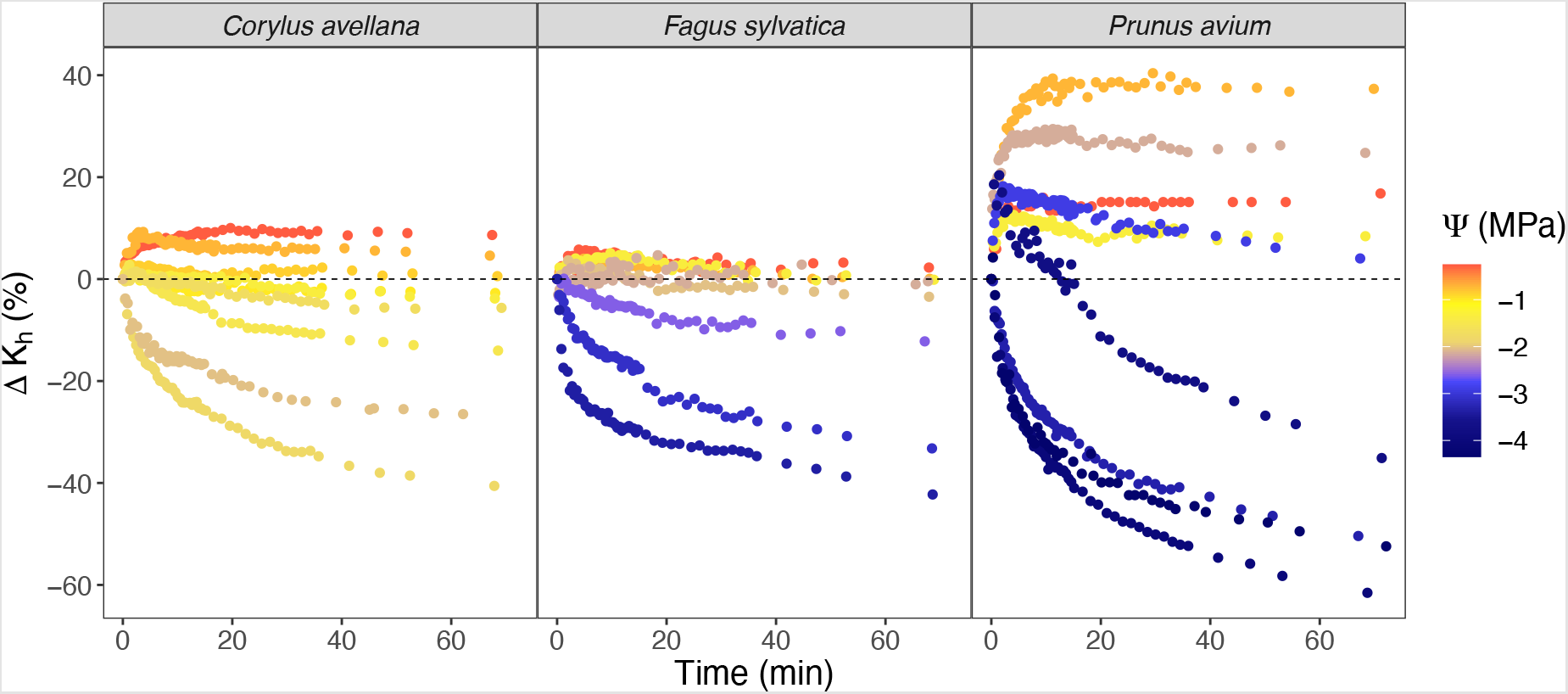
Changes in the hydraulic conductivity over time (ΔK_h_). Stem samples of three angiosperm species were spun in a flow-centrifuge for 1 hour at a constant temperature of 22°C and subject to a constant xylem water potential (Ψ) based on the rotational speed. ΔK_h_ is a relative value calculated through equation 2. Each colour represents a separate sample (n = 8 for *F. sylvatica* and n = 9 for *C. avellana*, and *P. avium*) that was spun at a fixed speed.

After one hour of spinning, ΔK_h_ decreased by a maximum of 40.6, 42.3, and 61.5%, and increased by a maximum of 10.0, 5.8, and 40.4% for *C. avellana, F. sylvatica* and *P. avium*, respectively. In general, we noticed that the magnitude of ΔK_h_ was variable, depending on the species and Ψ applied. Overall, the more negative the water potential induced in the flow-centrifuge, the higher the values of ΔK_h_. This interdependence of ΔK_h_ with time and Ψ was even more evident in our multiple non-linear regression of the relationship between ΔK_h_, time and Ψ, with R^2^ ranging from 0.69 to 0.94 (Fig. S3). In fact, Ψ did not affect the shape of any ΔK_h_ vs. time curve. Actually, Ψ affected the slope and the intercept coefficient of the regression. Although the three species studied showed overall similar patterns, they differed in the range of changes in K_h_, which affected the value of the coefficient in the regression.

Our results also showed a temperature effect on ΔK_h_, mainly at 5°C (Fig. 2). At this temperature and for Ψ values close to zero, we did not observe the same increasing pattern of ΔK_h_ as seen at 22°C. In fact, ΔK_h_ tended to increase during the first 5 minutes, similar to the pattern at 22°C, but then decreased and became negative after 15 minutes of spinning (except for one sample of *P. avium*), reaching values of -23, -24, and -8%, for *C. avellana, F. sylvatica*, and *P. avium*, respectively. Contrary, for more negative Ψ values at 5°C, ΔK_h_ presented a pattern similar to that observed at 22°C, decreasing by a maximum of -23, -33, and -67%, for *C. avellana, F. sylvatica* and *P. avium*, respectively. When spinning samples at 35°C, ΔK_h_ showed a similar trend over time to what was observed at 22°C, except for *P. avium*, which reached 78.5 and -67.4% as the most positive and negative values of ΔK_h_, respectively.

**Fig. 2.**
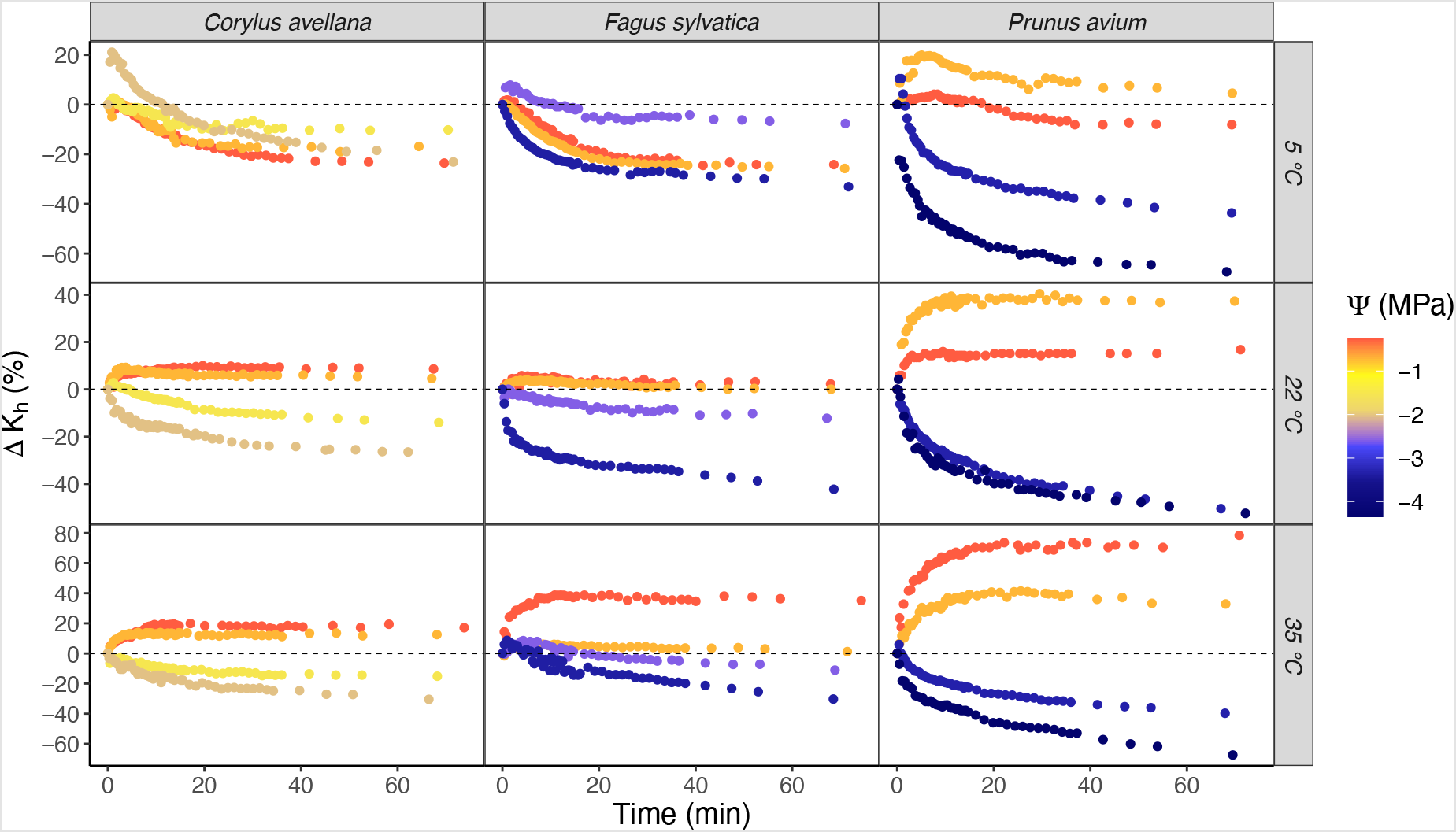
Changes in the hydraulic conductivity over time (K_h_) for stem samples of three angiosperm species (*C. avellana, F. sylvatica* and *P. avium*), which were spun in a flow-centrifuge for 1 hour. The two highest and two lowest rotational speeds applied at 22°C (Fig. 1) were also tested at 5° and 35°C. The samples were subjected to a constant xylem water potential (Ψ) based on the rotational speed applied. ΔK_h_ is a relative value calculated through equation 2. Each colour represents a separate sample that was spun at a fixed speed.

The rectangular hyperbola model demonstrated a significant prediction of K_h_ values (p-value < 0.05) when we considered the two most negative Ψ values applied (Fig S4 and S5). The low values of the Root mean square error (RMSE) showed that this model could predict K_h_ values with less than 1.3×10^−6^ % of error. Therefore, we could estimate the minimum value of hydraulic conductivity (K_min_), and consequently the relative hydraulic conductivity (K_relative_) with high precision for most of the treatments. However, the model underestimated K_min_ for *F. sylvatica* at 35 °C. In this particular case, the increase of K_h_ verified at the beginning of the experiment shifted the model estimation, resulting in lower values of K_min_ and K_relative_.

### Experiment 3: Location of embolised vessels in stem segments

The Percentage Loss of Conductivity (PLC) changed over time and was not homogeneously distributed along the 27.4 cm stem sample (Fig. 3). The distribution of the PLC was symmetrical in relation to the centre, except for *F. sylvatica* at 30 and 60 minutes, which presented higher values of PLC on the downstream. For all species studied, the PLC was significantly higher in segments at the centre of the centrifuge sample (p-value < 0.001) than in segments near the stem ends. The PLC in segments at the centre of the centrifuge sample increased with a longer spinning time (p-value < 0.05). This trend was most evident for *C. avellana*, with PLC increasing from 7.8% after 1 minute of spinning, to 53% after 7.5 minutes of spinning. *C. avellana* also showed high values of PLC at both stem ends, which were gradually decreasing while the amount of embolism increased over time near the centre of the centrifuge sample.

**Fig. 3.**
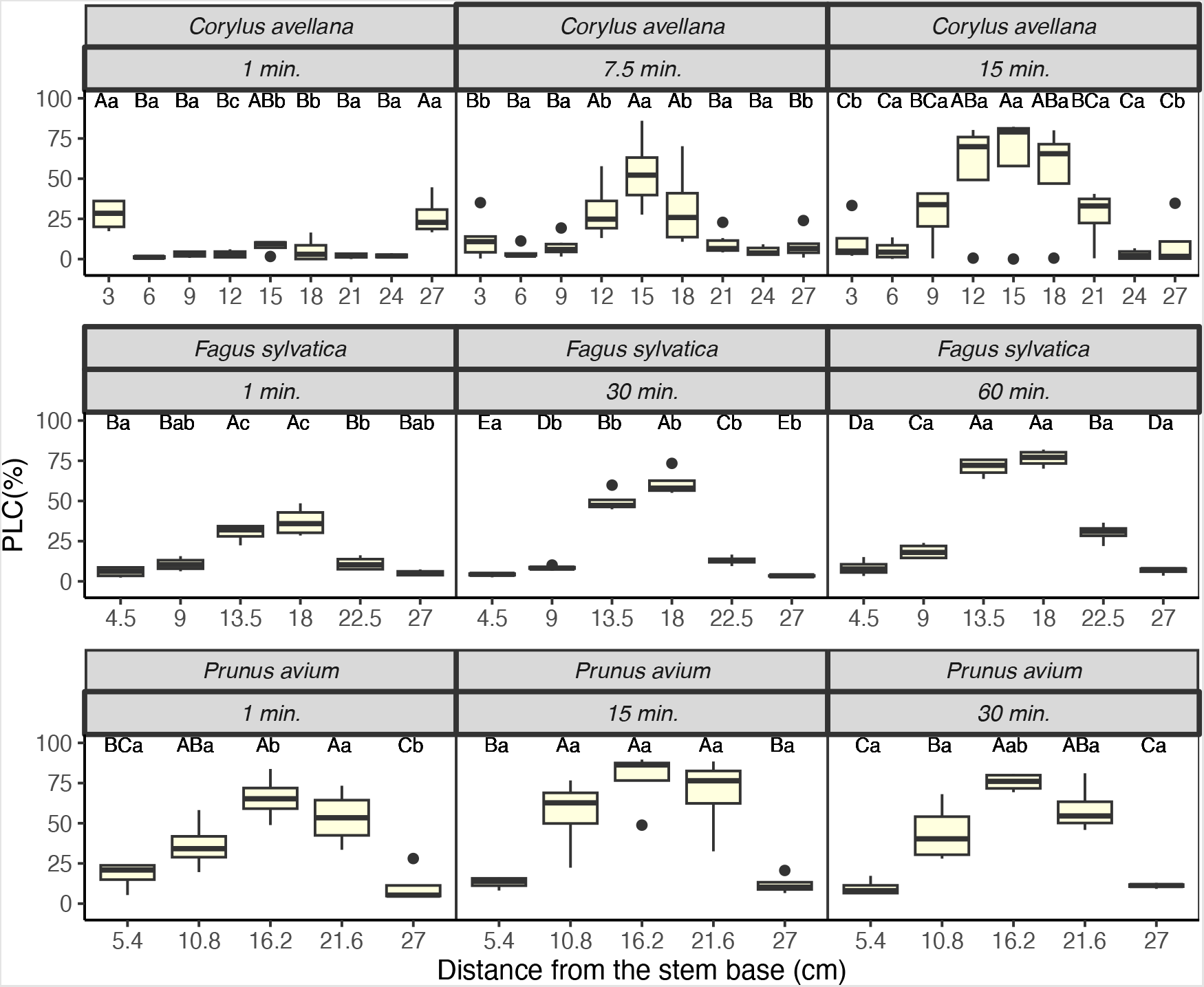
Percentage Loss of Conductivity (PLC) in 27.4 cm long stem samples of three angiosperm species. Stem samples were spun in a flow-centrifuge at a constant temperature (22°C) and water potential (−2.00, -3.35 and -4.35 MPa for *C. avellana, F. sylvatica* and *P. avium*, respectively) but different spinning times. After spinning, samples were re-cut in short segments and, then connected to an Ultra-Low Flowmeter to take hydraulic conductivity measurements. PLC was estimated for each of the segments using the measurements taken before and after flushing the samples. The spinning time and the segment location were the sources of variation. When significant differences between averages were observed, Tukey’s HSD tests (P < 0.05, and n = 4) were applied. Capital letters indicate significant differences among segments (same sample), and lowercase letters indicate significant differences among spinning times (same species).

### Modelling of the gas pressure in a recently embolised vessel in flow-centrifuge experiments

Once embolism had occurred near the centre of a centrifuge sample, the model predicted fast movement of gas along the 27.4 cm stem sample during the first minutes, but considerably slower gas diffusion rates after 1 hour (Fig. 4). The relative gas concentration in a recently embolised vessel (C_relative_) increased therefore quickly, reaching 50% after 2.8 to 8.0 minutes of the embolism event simulation based on the anatomical parameters of *C. avellana* and *P. avium*. This process was less pronounced and slowest for *F. sylvatica*, which reached C_relative_ of 50% after 16.6 to 31.8 minutes. One hour after embolism formation, C_relative_ reached values between 83.8 and 87.7, 63.1 and 69.7, and 79.6 and 84.5% for *C. avellana, F. sylvatica* and *P. avium*, respectively.

**Fig. 4.**
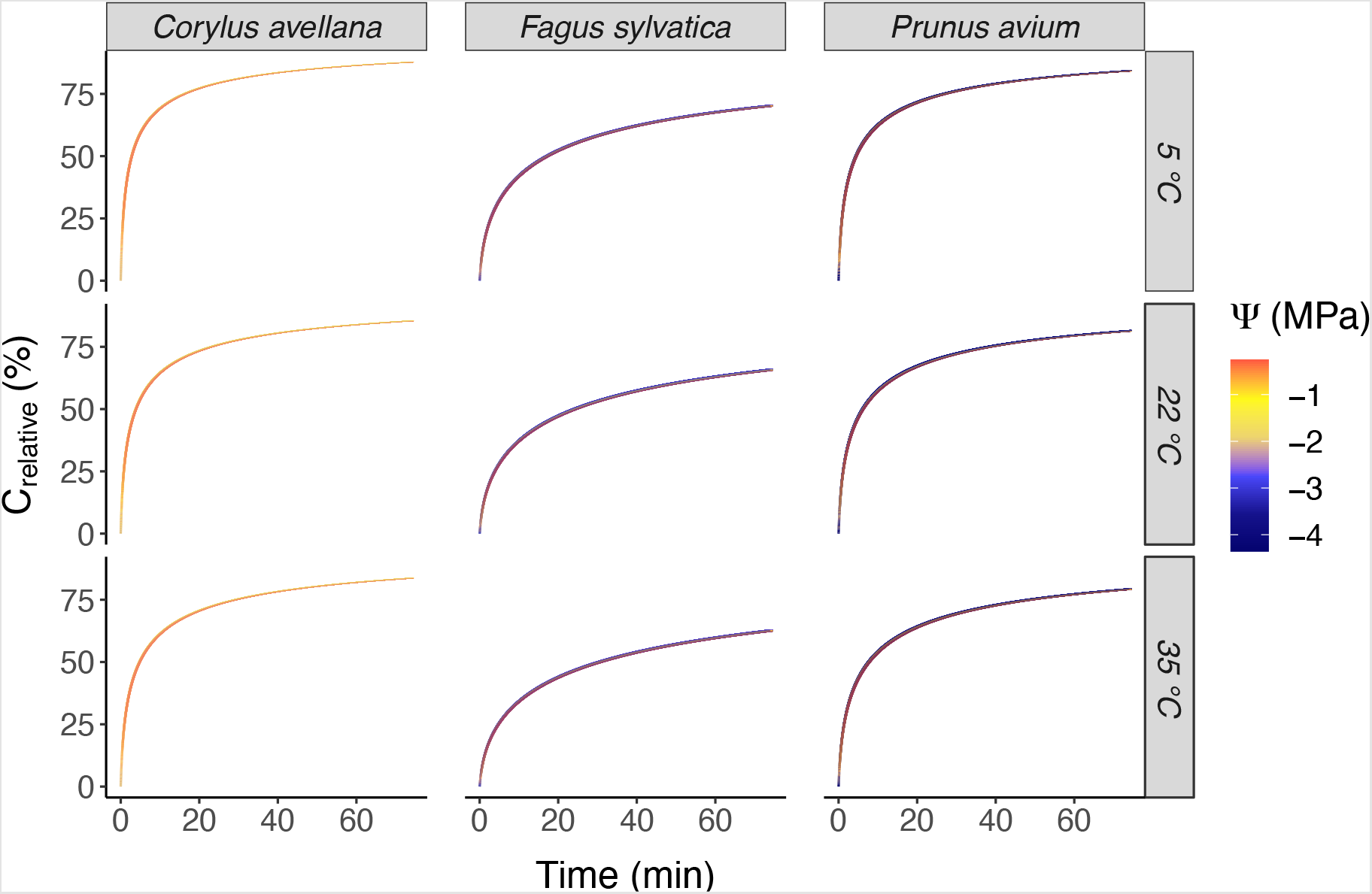
Temporal dynamics of the relative modelled gas concentration in a recently embolised vessel (C_relative_) in the centre of a centrifuge sample. After an embolism event, the simulation assessed the axial diffusion along the 27.4 cm stem samples while it was centrifuged. The model was based on the Unit Pipe Pneumatic model of Yang *et al*. (2023) and considered the same conditions of temperature and water potential (Ψ) applied to flow-centrifuge measurements. C_relative_ is a relative value, which was normalized by equation 21 considering its lowest (vapour-filled vessel) and highest (atmospheric concentration) value.

The axial gas diffusion rate was strongly affected by the temperature and the anatomical traits of the species, but minimally affected by Ψ. The higher the temperature, the smaller the changes in C_relative_ over time for all three species (Fig. S6). In contrast to temperature, Ψ tended to accelerate the gas diffusion kinetics. However, its effect was small, changing C_relative_ less than 1%. Between species, we noticed a slight difference between *C. avellana* and *P. avium*, but a larger difference between these species and *F. sylvatica*. (Fig. S7). For *F. sylvatica*, the axial gas diffusion was on average 20% slower than for *C. avellana* and *P. avium*.

We observed a strong and positive correlation between the relative hydraulic conductivity (K_relative_) and our modelled values of the gas concentration in a recently embolised vessel in the centre of a centrifuge sample (C_relative_) (Fig. 5). These correlations were significant (p < 0.05) for all conditions of temperature and Ψ. We also observed low values of RMSE and a Person correlation coefficient (R) close to 1. Moreover, both RMSE and R changed among the species. The performance of the model was most accurate for *P. avium*. In this species, the model predicted the measured data with an error (RMSE) up to 6.8% for a Ψ of -3.29 MPa and a temperature of 22°C. However, when considering *F. sylvatica* at 35°C, our model underestimated K_min_, and consequently K_relative_, resulting in RMSE values of 35.5 and 28.1% for Ψ of -2.57 and -3.35 MPa, respectively. For *C. avellana* at 5° and 22°C, the loss of K_h_ was slower than the increase in C_relative_. On the other hand, *F. sylvatica* at 22°C and at a Ψ of -4.35 MPa showed an underestimation of C_relative_, with K_h_ decreasing faster than the increasing C_relative_. For *C. avellana* at 35°C, *F. sylvatica* at 5°C and *P. avium* at 5, 22, 35°C, the correlation curves were close to the 1:1 line (Fig. 5), showing that the increase in C_relative_ was strongly correlated to the decrease in K_relative_ with a maximum error of 21.1%.

**Fig. 5.**
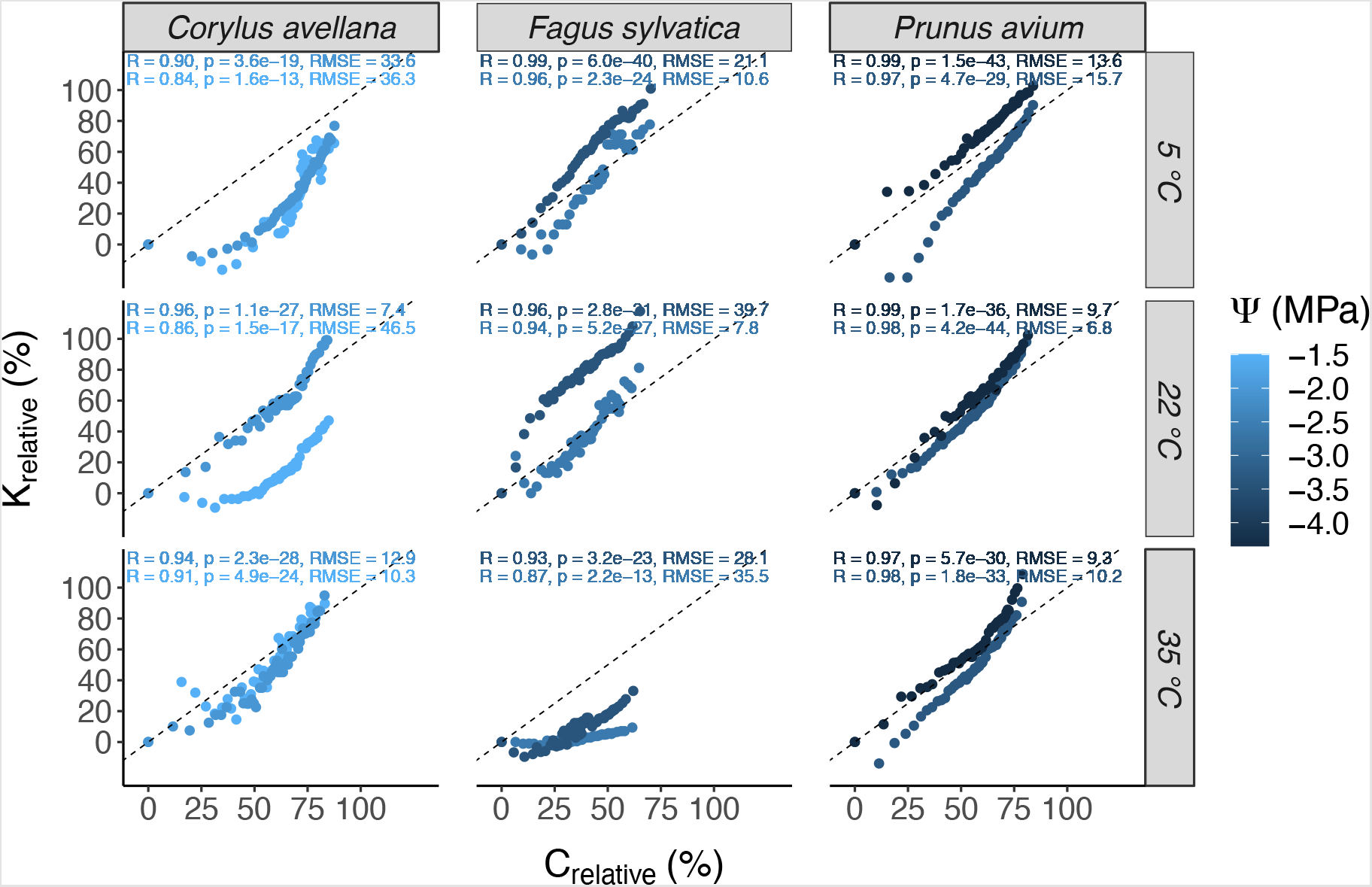
Correlation between modelled gas concentration in a recently embolised vessel (C_relative_) in the centre of a centrifuge sample, and relative hydraulic conductivity (K_relative_) for the three species studied (*C. avellana, F. sylvatica* and *P. avium*), at three temperatures (5, 22 and 35°C), and the two most negative water potentials (Ψ) applied in experiment 2. K_relative_ is a relative value of the hydraulic conductivity, which was normalized by equation 3, considering the first hydraulic conductivity measurement and the minimum value predicted by the rectangular hyperbola model. Each colour represents a correlation between C_relative_ and K_relative_ with its respective Person correlation coefficient (R), root mean square error (RMSE) and p-value (p). The black dashed line is a 1:1 line, which represents our hypothesis.

## Discussion

Our results revealed that hydraulic conductivity (K_h_) measurements in a flow-centrifuge are not stable at a fixed pressure and temperature over a spinning duration of 1 hour. Since relative changes in hydraulic conductivity (ΔK_h_) did not change randomly around zero over time, the variation observed cannot be considered as noise (Fig. 1 and 2). As expected, experiment 2 (the effect of water potential, time and temperature on K_h_) showed that pressure is highly important in determining the overall magnitude of K_h_, with absolute K_h_ values taken at high water potentials being much higher than those measured at low water potentials (Fig. S2). However, depending on the temperature, the xylem anatomy of the plant species studied, and the pressure applied, changes in K_h_ showed a particular behaviour over time, increasing or decreasing at a particular rotational speed. Therefore, if decreases in K_h_ are at least partly related to increasing levels of embolism, we argue that embolism spread may represent a relatively slow process, and that embolism is not only pressure-driven, but also affected by time. The main reason for using ΔK_h_ is that we aim to shed light on the relative changes in K_h_ over time to explore the driving mechanisms behind embolism spread. These ΔK_h_ values also enable us to compare the effects of the different treatments applied, and differences in xylem anatomy between species (Fig. 1, 2 and S3). As such, methodological issues related to vulnerability curves and how we quantify embolism resistance go beyond the purpose of this study, but will be discussed in future work. Our findings presented strong evidence for a mechanistic relationship between K_h_ and embolism formation. While an increase in the Percentage Loss of Conductivity (PLC) over time at the centre of centrifuge samples (Fig. 3) can explain the decrease in K_h_ in flow centrifuge measurements (Fig. 1 and 2), experiment 1 (the effect of flow on K_h_) showed that temporal variation in K_h_ cannot be driven only by a non-embolism related decline in K_h_ over time. We observed a small, but long-term decline in conductivity when there was a constant flow through a stem sample (Fig. S1), which can be attributed to various processes (Sperry *et al*., 1988; Yin *et al*., 2019). Examples of potential explanations include pit membrane clogging, wound response, a potential difference between the inlet and outlet flow, radial or background flow, or compression of intervessel pit membranes (Espino & Schenk, 2011; De Baerdemaeker *et al*., 2019). However, K_h_ was minimally affected over time when we injected water only during the actual K_h_ measurements, with K_h_ changing by 0.7% within the first spinning hour. This “natural” decline in K_h_ induced by flow cannot explain the much higher changes in K_h_ observed in experiment 2, and is certainly not in line with our observations of increases in ΔK_h_ by up to 32.8% and even 78.5%. Moreover, if tiny gas bubbles included in the injection water of the centrifuge would cause embolism, as suggested by Yin *et al*. (2019), we would only observe a decrease in K_h_ and higher levels of embolism at the injection side, which was not the case.

Since embolism removal is generally not considered to occur in a sample that is exposed to a negative pressure (Tyree & Sperry, 1988; Lamarque *et al*., 2018), the observed increase in K_h_ over time has been rarely reported in literature. Sperry *et al*. (1988) described this increase as a rearrangement and/or dissolution of emboli within xylem conduits, which mainly occurs under subatmospheric pressure (6 KPa). Thus, this rearrangement and/or dissolution of emboli can explain the increase in K_h_ when we applied a water potential close to zero as well as at the beginning of slightly negative pressures. Apart from rearrangement, the replacement of xylem sap with the flow-centrifuge solution may modify the water permeability of interconduit pit membranes and thus increase K_h_, a phenomenon also known as the ‘ionic effect’ (Jansen *et al*., 2011). Interestingly, samples centrifuged at 5°C presented a much weaker increase in ΔK_h_ (Fig 2), suggesting that the temperature may also play an important role in the rearrangement and/or dissolution of emboli and eventually in embolism spreading. One explanation for this finding can be provided by Briggs (1950), who described that water under negative pressure loses much of its ability to support negative pressure between 0 and 5°C, which may lead to an increase in embolism events. While the likelihood of homogeneous cavitation is known to increase with increasing temperature (Caupin & Herbert, 2006; Herbert *et al*., 2006), embolism in xylem conduits is very likely not caused by homogeneous cavitation (Hölttä *et al*., 2002). Overall, little is known about how temperature may affect xylem embolism resistance (Cochard *et al*., 2007; Lodge *et al*., 2018). Temperature may also affect gas solubility and oversaturation of xylem sap, or could affect the dynamic surface tension at gas-liquid interfaces (Schenk *et al*., 2016; Yang *et al*., 2020).

Experiment 3 (embolism location) shows that there are at least two different locations in centrifuge samples where embolism is formed, namely near the stem edges and in the centre. Over time, the amount of embolism near stem edges decreased, but increased near the centre. We speculate that the initial embolism levels near stem edges are caused by the axial proximity of the cut-open conduits to atmospheric gas, either dissolved gas in the liquid in the cuvettes, or gas outside the cuvettes. However, additional experiments would be needed to test this hypothesis. The most negative pressure in the centre clearly increases the likelihood of new embolism events in this area, which is associated with a constant decrease in K_h_ over time. It is known that gas solubility increases slightly with decreasing negative liquid pressure (Mercury *et al*., 2003; Lidon *et al*., 2018), but the effect of temperature has a higher influence on gas solubility than pressure (Schenk *et al*., 2016). Our results also revealed that the speed of embolism formation and the location of embolism may vary according to the species studied. Therefore, we suggest that the disagreement between earlier studies on the location of embolised vessels in centrifuged stems is likely due to different species used in each study, as well as to the spinning time, which proved to be essential in our observations, but was not reported or considered in earlier work (Cai *et al*., 2010; Cochard *et al*., 2010; Sperry *et al*., 2012; Tobin *et al*., 2013; Martin-StPaul *et al*., 2014; Yin *et al*., 2019).

The difference in the speed of embolism propagation between species is related to the ability of the species to resist embolism formation, and is likely associated with the vessel anatomy, including vessel diameter and length, and intervessel connectivity. However, more experiments with a larger number of species would be needed to test this hypothesis (Lens *et al*., 2011, 2022; Isasa *et al*., 2023). Interestingly, the vessel anatomy also affected our model of gas pressure in a recently embolised vessel in the centre of a centrifuge sample. While a higher value of the total intervessel pit membrane area (*A*_p_) increases the axial gas transport rate in the sap and across the pit membranes (k_a_ and k_p_, respectively), a larger vessel volume (*V*_v_) requires much more gas and probably more time to achieve atmospheric pressure. Wide and long vessels can therefore be suggested to provide a longer, temporary “buffer” against further embolism spread than narrow and short ones. If the gas concentration in a recently embolised vessel (C_relative_) is related to the rate of embolism formation and propagation, it seems reasonable to expect that *V*_v_ and *A*_p_ could play a role in xylem embolism resistance.

Although it is well known that pressure triggers embolism formation, the mechanisms behind embolism initiation are largely unsolved (Schenk *et al*., 2015). Here, we modelled the transition of a water-vapoured filled conduit to a conduit filled with gas at atmospheric pressure, which was shown to correlate significantly with ΔK_h_. This significant correlation provides strong evidence that the increase in C_relative_ and the decline of ΔK_h_ have the same time scale and order of magnitude, suggesting that a functional link between both processes is likely. Two different steps in the process of embolism spread may explain this functional link. Firstly, a recently embolised vessel that is water vapour filled provides likely a buffer for further embolism propagation because the low vapour pressure (ca. 3.2 kPa) will extract gas molecules from surrounding vessels, until it eventually reaches atmospheric pressure (Fig. 6; Wang *et al*., 2015). If a high gas concentration is necessary for embolism initiation, the temporal and local extraction of gas from surrounding, sap-filled vessels can be suggested to reduce the risk of new embolism events. Secondly, when sufficient gas has been accumulated in an embolised conduit, this gas reservoir becomes a gas source that may increase the likelihood to have embolism formation in a neighbouring, interconnected conduit (Guan *et al*., 2021). Thus, we assume that the amount of gas in a recently embolised conduit increases the nucleation process of embolism occurring in a neighbouring conduit. Finally, this new conduit will again extract gas from neighbouring conduits, slowing down embolism spread.

**Fig. 6.**
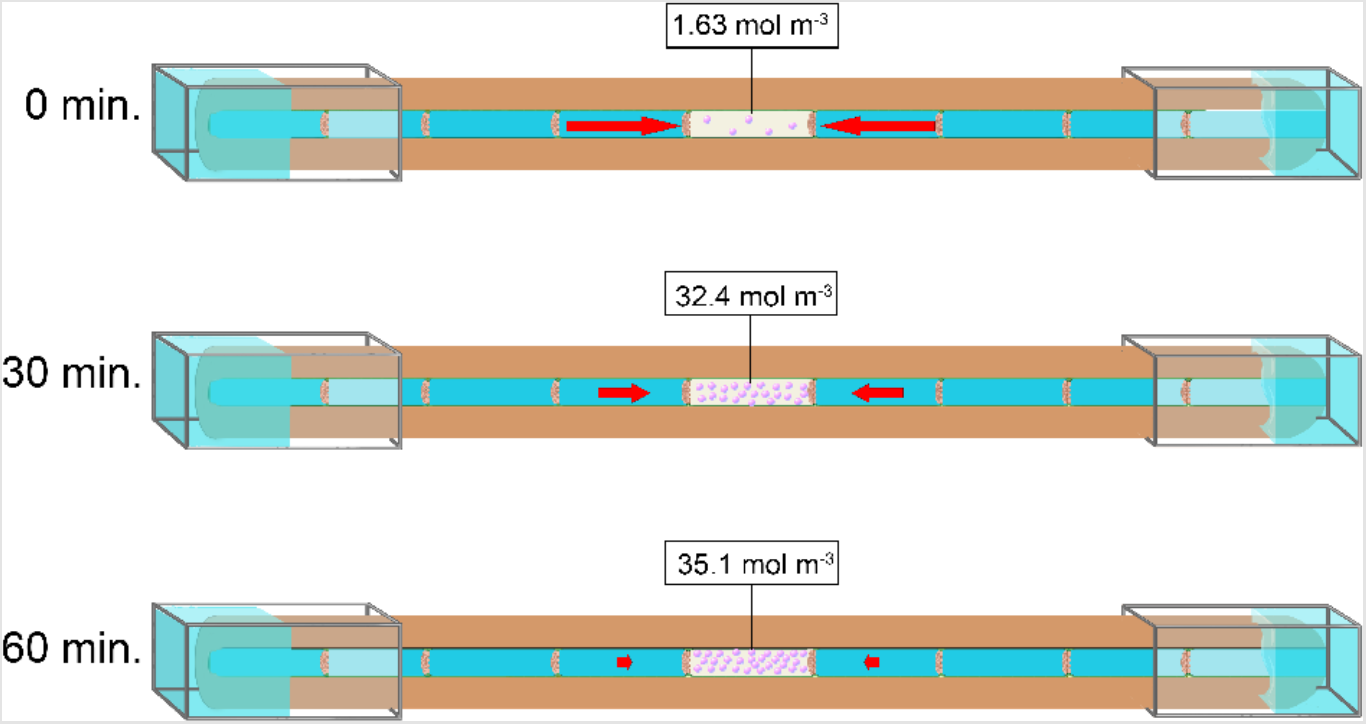
Illustration of dynamic changes in the modelled gas pressure in a recently embolised vessel in the centre of a centrifuge sample. The figure presents the absolute values of the modelled gas concentration in a recently embolised vessel at three different time steps (0, 30, and 60 min.) for *Corylus avellana* at a temperature of 22°C and a water potential of -2.00 MPa.

We cannot exclude that minor changes in K_h_ may not always reflect changes in the amount of embolism, which is indirectly estimated based on K_h_. This fact can propagate a certain degree of error in our measurements, and may weaken our analyses, correlations, and interpretation of the measurements. Nevertheless, adjusting the data measured in models proved to be an appropriate strategy as this approach ensured that the data used in our analyses were not randomly generated but the result of physical phenomena. Hence, experiments with flow-centrifuges provide a valid approach to investigate the driving mechanisms behind embolism, even if flow-centrifuge experiments do not entirely copy functional xylem operating under natural field conditions in intact plants.

In conclusion, our experiments showed that xylem pressure is the most important determinant in initiating a given level of embolism in xylem conduits. However, embolism propagation proved to be dynamic, with pressure, time, and temperature playing an interdependent role in embolism spread. Not only the amount of embolism may slightly change over time in a centrifuge, but also its location within centrifuge samples. Our experimental and modelled data provide evidence that embolism propagation relies on gas movement, which is driven by local differences in gas concentration. Moreover, embolism propagation is shown to be related to vessel anatomy. A high total intervessel pit membrane area in a large vessel increases the axial gas transport rate, but a large, recently embolised vessel may also require more gas and therefore more time to achieve atmospheric pressure than a narrow recently embolised vessel. Overall, our findings raise questions about the traditional assumption that embolism is exclusively pressure-driven, as suggested for instance by the x-axis of vulnerability curves. Our results also contribute to our understanding that embolism resistance of plant xylem represents a relative and dynamic trait that is affected by multiple parameters.

## Acknowledgements

We thank Mathias Eberhardt and Merlin Grimm for experimental work with the ChinaTron centrifuge. The authors also thank the valuable insight provided by Jochen Schenk.

## Abbreviations

ΔK_h_: Relative change in hydraulic conductivity
C_relative_: Relative modelled gas concentration in a recently embolised vessel
K_0_: Initial hydraulic conductivity
K_0_’: Hydraulic conductivity before flushing a stem segment
K_F_’: Hydraulic conductivity after flushing a stem segment
K_h_: Current hydraulic conductivity
K_min_: minimum hydraulic conductivity
K_relative_: relative hydraulic conductivity
PLC: Percentage Loss of Conductivity
R: Person correlation coefficient
RMSE: root mean squared error
Ψ: Minimum xylem water potential in the middle of a stem sample

## Supporting Information

**Fig. S1** Relative change in hydraulic conductivity over time (ΔK_h_, %).

**Fig. S2** Exact values of hydraulic conductivity (K_h_) measured over time.

**Fig. S3** Relative changes in hydraulic conductivity (ΔK_h_) as a function of time and water potential.

**Fig. S4** Measured vs. predicted hydraulic conductivity (K_h_) values over time for three species studied (*C. avellana, F. sylvatica* and *P. avium*), three temperatures (5°, 22° and 35°C) and three water potentials.

**Fig. S5** Measured vs. predicted hydraulic conductivity (K_h_) values over time for three species studied (*C. avellana, F. sylvatica* and *P. avium*), three temperatures (5°, 22° and 35°C) and three water potentials.

**Fig. S6** Temporal dynamics of the relative gas concentration modelled in a recently embolised vessel (C_relative_) for the three species studied (*C. avellana, F. sylvatica* and *P. avium*) and three temperatures applied (5 °, 22 ° and 35 °C).

**Fig. S7** Temporal dynamics of the relative gas concentration modelled in a recently embolised vessel in the centre of a centrifuge sample (C_relative_) for three species (*C. avellana, F. sylvatica* and *P. avium*).

**Table S1** Temperature, rotational speed and water potential applied in Experiment 2 to three angiosperm species studied with a flow-centrifuge.

**Table S2** Overview of the stem segments used to assess the distribution of embolised vessels in experiment 3.

**Table S3** Characteristics of the xylem vessels of the three species studied.

**Table S4** Intercept coefficient (*a*), water potential coefficient (*b*), and temperature coefficient (*c*) for Henry’s constant (K_H_) calculation of different gases (equation 8).

**Table S5** Values of several variables that were required to model the gas pressure in a recently embolised vessel in flow-centrifuge experiments.

**Table S6** Overview of all abbreviations, units, definitions and equations used in the gas diffusion model for flow-centrifuge measurements.

**Methods S1** Modelling of the gas pressure in a recently embolised vessel in flow-centrifuge experiments.

**Methods S2** Multiple regression for ΔK_h_ as a function of time and Ψ.

**Methods S3** Fitting a rectangular hyperbola model.

**Box S1** Workflow of the axial gas diffusion model for flow-centrifuge experiments.

