## Supporting Information for "Gas diffusion kinetics drive embolism spread in angiosperm xylem: evidence from flow-centrifuge experiments and modelling"

**Fig. S1** Relative change in hydraulic conductivity over time ($\Delta K_h$, %) in stem samples of *C. avellana* spun in a flow-centrifuge for 8 hours at a constant temperature of 22 °C and subjected to a constant xylem water potential of -1.17 MPa. $\Delta K_h$ is a relative value calculated by equation 2. Each colour represents a separate sample that was spun under two different conditions: (●) continuous addition of flow solution into the cuvettes, resulting in continuous flow through the sample, and (●) no addition of solution between hydraulic conductivity measurements, with flow across the sample limited to short time-interval when hydraulic conductivity was measured.

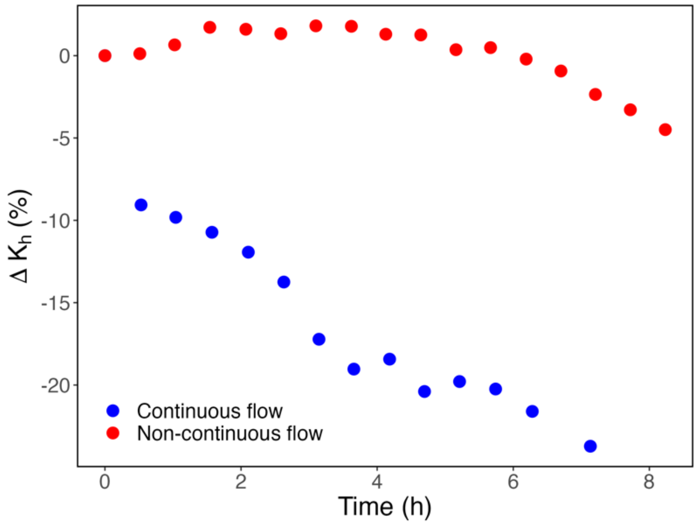

**Fig. S2** Exact values of hydraulic conductivity ($K_h$) as measured over time in stem samples of three angiosperm species, which were spun in a flow-centrifuge for 1 hour at a constant temperature of 22 °C, and subject to a constant xylem water potential (Ψ) based on the rotational speed chosen. Each colour represents a separate sample (n = 8 for *F. sylvatica* and n = 9 for *C. avellana*, and *P. avium*) that was spun at a fixed speed.

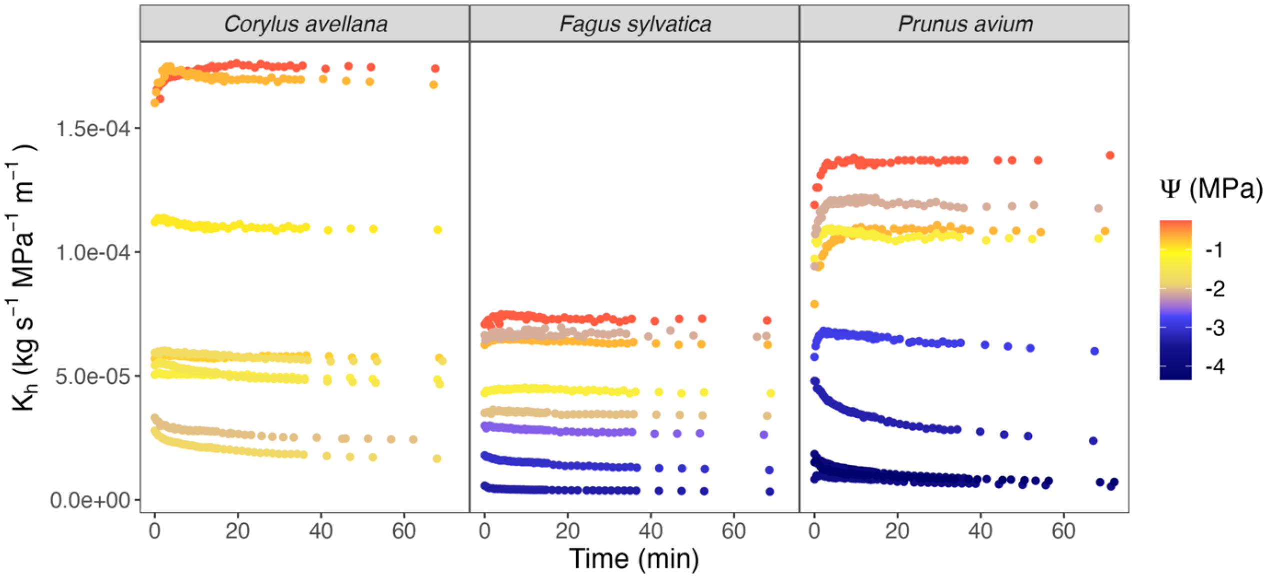

**Fig. S3** Relative changes in hydraulic conductivity ($\Delta K_h$) as a function of time and water potential ($\Psi$). Stem samples of three angiosperm species were spun in a flow-centrifuge for 1 hour at a constant temperature of 22 °C, and subject to a constant xylem water potential ($\Psi$) based on a given rotational speed. Each curve consisting of multiple black dots represents a separate sample (n = 8 for *F. sylvatica* [b] and n = 9 for *C. avellana* [a], and *P. avium* [c]), which were spun at a fixed speed. $\Delta K_h$ is a relative value calculated by equation 2. The colourful surfaces represent simulated data from the multiple non-regression equations shown at the top of each graph with their corresponding $R^2$.

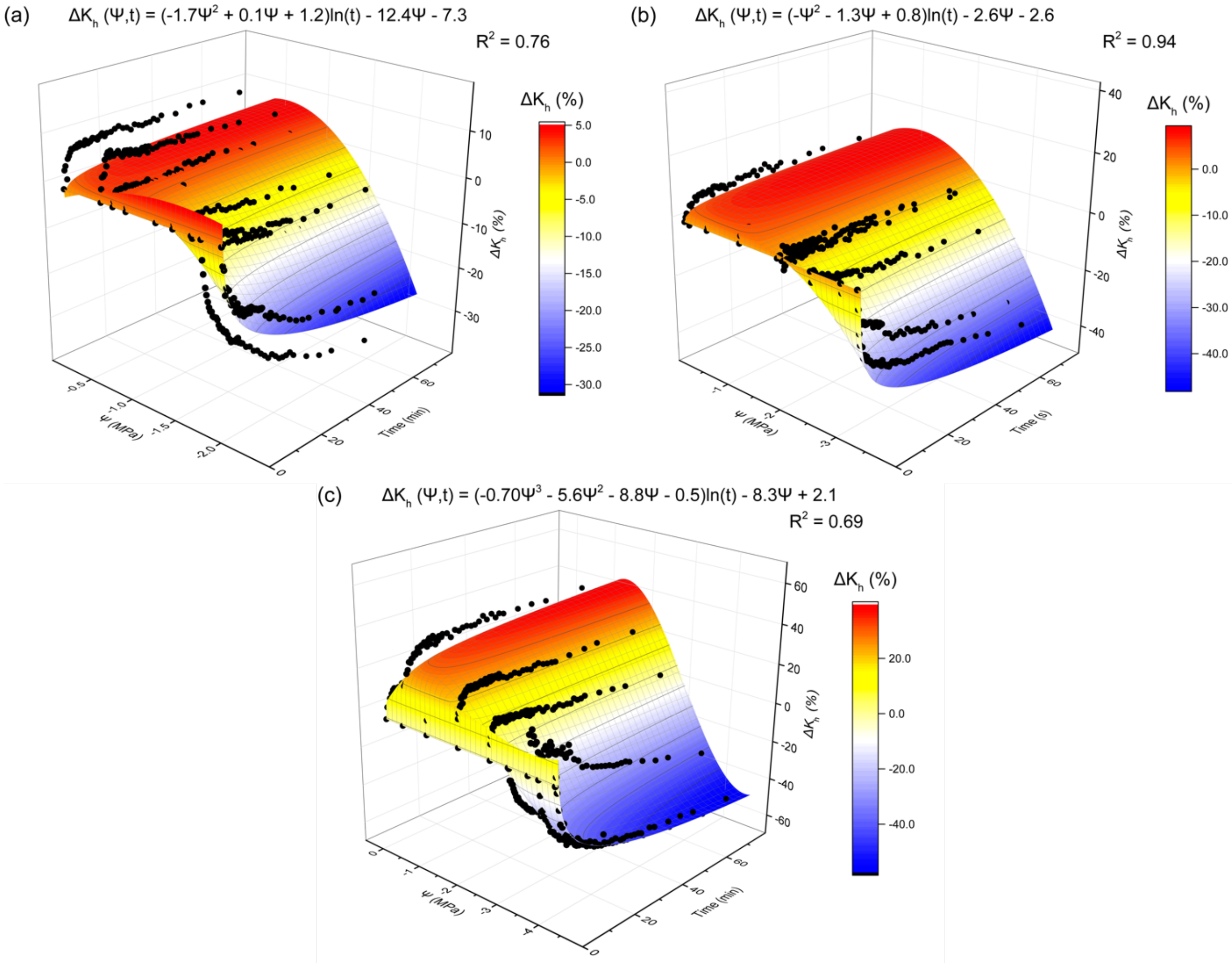

**Fig. S4** Measured (dots) vs. predicted (full line) hydraulic conductivity ($K_h$) values over time for three angiosperm species (*C. avellana*, *F. sylvatica* and *P. avium*) at three different temperatures (5 º, 22 º and 35 ºC) and at three water potentials (Ψ). Dots represent stem samples that were spun in a flow-centrifuge for 1 hour at a constant temperature and subject to a constant Ψ based on a given rotational speed. The curves are predicted values of $K_{h,}$, which were estimated by a rectangular hyperbola model. The minimum value of hydraulic conductivity ($K_{min}$) was estimated using the model equation provided. The root mean squared error (RMSE), Person correlation coefficient (R), and p-value (p) were used as a measure of the goodness of fit.

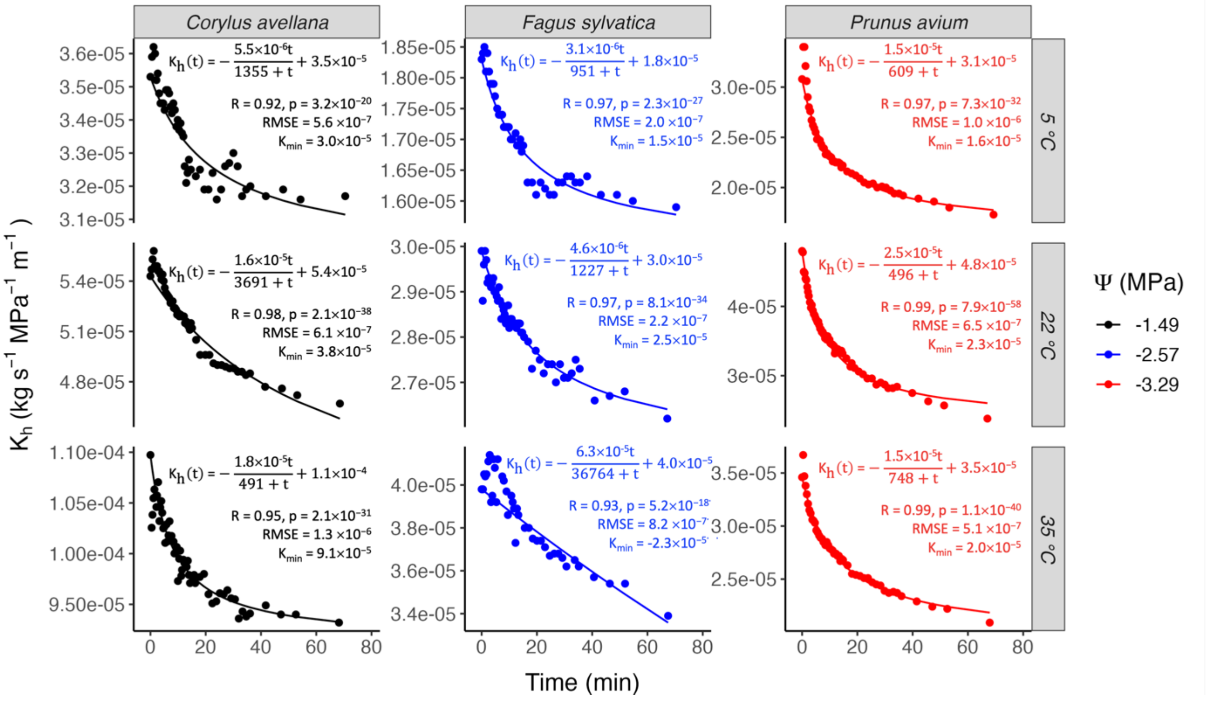

**Fig. S5** Measured (dots) vs. predicted (full line) hydraulic conductivity ($K_h$) values over time for three angiosperm species (*C. avellana*, *F. sylvatica* and *P. avium*), at three different temperatures (5º, 22º and 35ºC) and three water potentials (Ψ). Dots represent stem samples that were spun in a flow-centrifuge for 1 hour at a constant temperature and subject to a constant Ψ based on a given rotational speed. The curves are predicted values of $K_h$, which were estimated by a rectangular hyperbola model. The minimum value of hydraulic conductivity ($K_{min}$) was estimated using the model equation provided. The root mean squared error (RMSE), Person correlation coefficient (R) and p-value (p) were used as a measure of the goodness of fit.

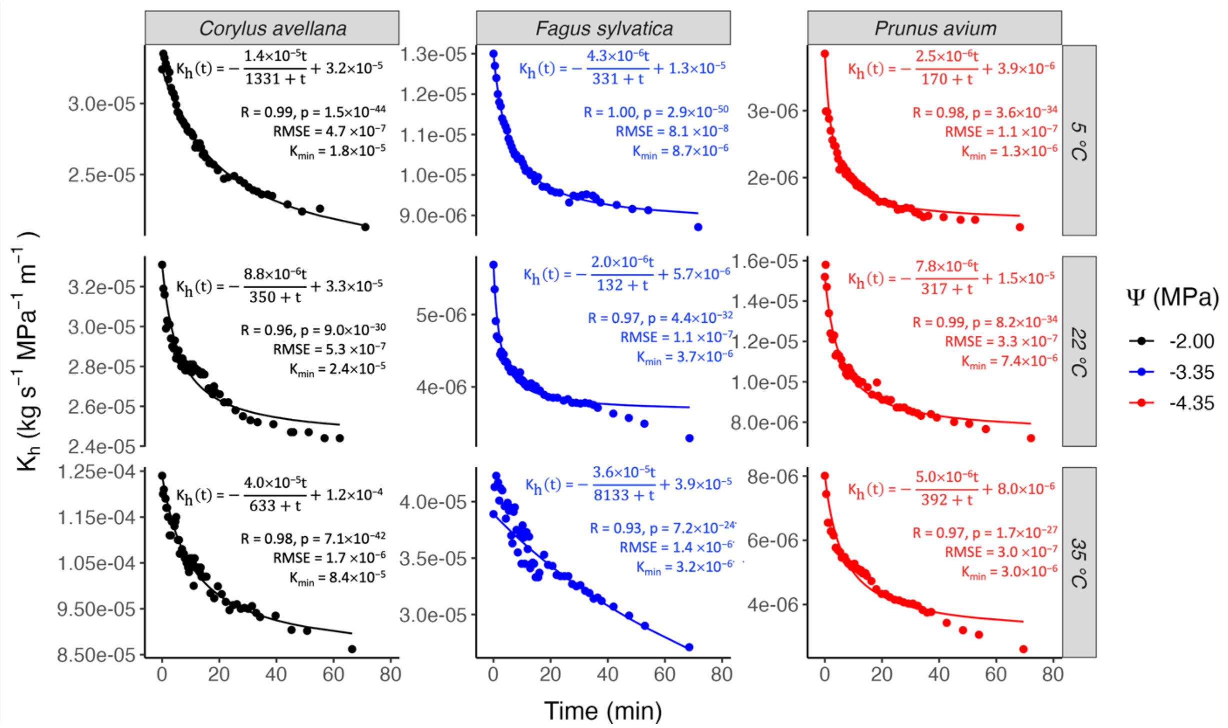

**Fig. S6** Temporal dynamics of the relative gas concentration modelled in a recently embolised vessel ($C_{relative}$) in the centre of a centrifuge sample. Shown are data for the three species studied (*C. avellana*, *F. sylvatica* and *P. avium*) at three temperatures (5°, 22° and 35°C). After an embolism event, the simulation assessed the axial diffusion of gas along the 27.4 cm stem samples during centrifugation. The model was based on the Unit Pipe Pneumatic model of Yang *et al.* (2023), and considered the conditions applied for flow-centrifuge measurements as well as the anatomical traits of the species studied. Data were retrieved from Fig. 4. In order to highlight the differences between species and the three temperatures, the various water potentials induced in our centrifuge experiments were combined. $C_{relative}$ was a relative value, which was normalized by equation 21 considering its lowest (100% water vapour-filled vessel) and highest (atmospheric gas concentration) value.

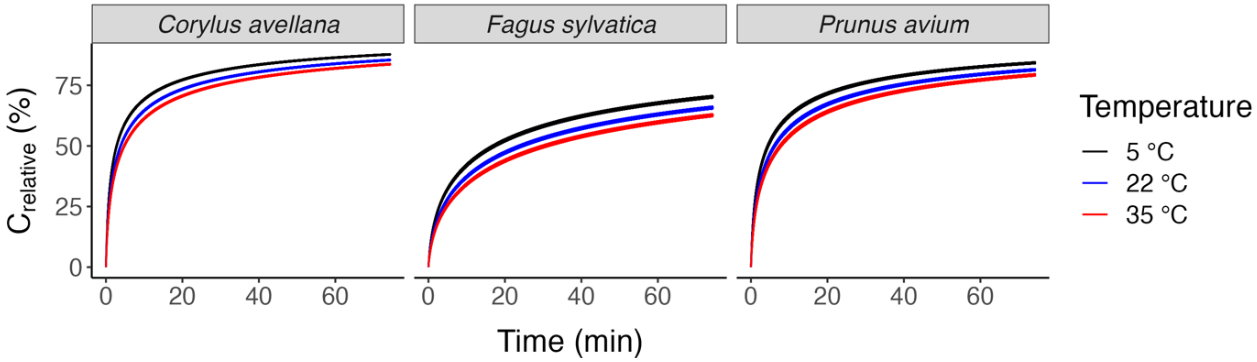

**Fig. S7** Temporal dynamics of the relative gas concentration modelled in a recently embolised vessel in the centre of a centrifuge sample ($C_{relative}$) for three species (*C. avellana*, *F. sylvatica* and *P. avium*). After an embolism event, the simulation assessed the axial diffusion along the 27.4 cm stem samples while these were centrifuged. The model was based on the Unit Pipe Pneumatic model of Yang *et al.* (2023) and considered the conditions applied for flow-centrifuge measurements as well as the anatomical traits of the species studied. Data retrieved from Fig. 4. In order to highlight the differences between species, the three temperatures and the water potentials applied to our centrifuge experiments were combined. $C_{relative}$ is a relative value, which was normalized by equation 21 considering its lowest (100% water vapour-filled vessel) and highest (atmospheric gas concentration) value.

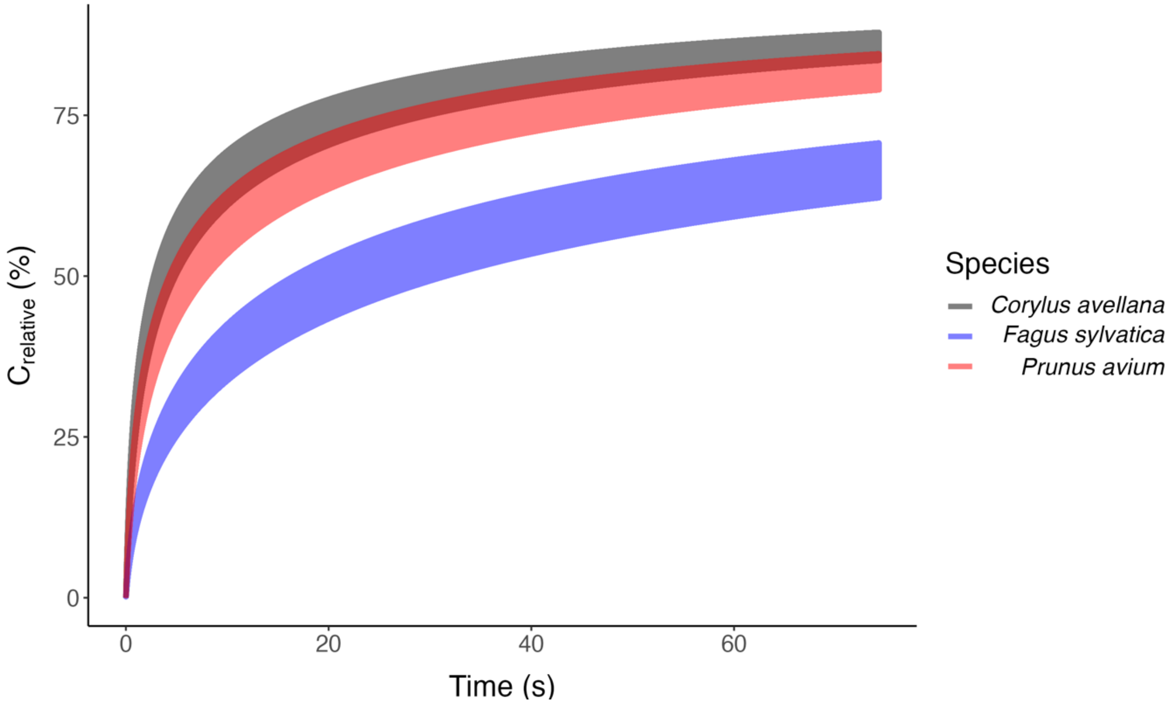

**Table S1** Temperature, rotational speed, and water potential applied in Experiment 2 to three angiosperm species studied with a flow-centrifuge.

| | *Corylus avellana* | | *Fagus sylvatica* | | *Prunus avium* | |
|---|---|---|---|---|---|---|
| Temperature | Rotational speed (RPM) | Water potential (MPa) | Rotational speed (RPM) | Water potential (MPa) | Rotational speed (RPM) | Water potential (MPa) |
| 5 °C | 2000 | -0.25 | 2000 | -0.25 | 2000 | -0.25 |
| | 3000 | -0.69 | 3000 | -0.69 | 3000 | -0.69 |
| | 4250 | -1.49 | 5500 | -2.57 | 6200 | -3.29 |
| | 4875 | -2.00 | 6250 | -3.35 | 7100 | -4.35 |
| 22 °C | 2000 | -0.25 | 2000 | -0.25 | 2000 | -0.25 |
| | 3000 | -0.69 | 3000 | -0.69 | 3000 | -0.69 |
| | 3250 | -0.83 | 4000 | -1.31 | 4000 | -1.31 |
| | 3500 | -0.98 | 4875 | -2.00 | 5000 | -2.11 |
| | 4000 | -1.31 | 5000 | -2.11 | 5800 | -2.87 |
| | 4250 | -1.49 | 5500 | -2.57 | 6200 | -3.29 |
| | 4500 | -1.69 | 6000 | -3.08 | 6600 | -3.74 |
| | 4625 | -1.79 | 6250 | -3.35 | 6800 | -3.98 |
| | 4875 | -2.00 | - | - | 7100 | -4.35 |
| 35 °C | 2000 | -0.25 | 2000 | -0.25 | 2000 | -0.25 |
| | 3000 | -0.69 | 3000 | -0.69 | 3000 | -0.69 |
| | 4250 | -1.49 | 5500 | -2.57 | 6200 | -3.29 |
| | 4875 | -2.00 | 6250 | -3.35 | 7100 | -4.35 |

**Table S2** Overview of the stem segments used to assess the distribution of embolised vessels in experiment 3. The segment length, number of segments, rotational speed and water potential varied between the species according to their vessel length and embolism resistance, which were based on Guan *et al.* (2022). The samples were subject to a temperature of 22°C and three different spinning times, which were based on preliminary trials (data not shown).

| Species | Segment length (cm) | Number of segments | Rotational speed (RPM) | Water potential (MPa) | Spinning time (min) |
|---|---|---|---|---|---|
| *Corylus avellana* | 3.0 | 9 | 4875 | -2.00 | 1 |
| | | | | | 7.5 |
| | | | | | 15 |
| *Fagus sylvatica* | 4.5 | 6 | 6250 | -3.35 | 1 |
| | | | | | 30 |
| | | | | | 60 |
| *Prunus avium* | 5.4 | 5 | 7100 | -4.35 | 1 |
| | | | | | 15 |
| | | | | | 30 |

**Table S3** Characteristics of the xylem vessels of the three species studied. Values indicate the mean and standard deviation of the equivalent circle diameter of vessels ($D$), mean vessel length ($L_v$), thickness of an intervessel pit membrane at the centre ($T_{PM}$), fraction of vessel wall in contact with other vessels ($F_c$), fraction of intervessel wall area occupied by intervessel pits ($F_{PF}$), fraction of total vessel wall area occupied by intervessel pits ($F_P$), total pit membrane surface area ($A_p$), and vessel volume ($V_v$). Values of $D$, $L_v$, $T_{PM\_centre}$, $F_C$, and $F_{PF}$ were based on Guan *et al.* (2021). The estimation of $F_P$ and $A_p$ followed the model of Wheeler *et al.* (2005). The calculation of $V_v$ considered a vessel as a cylinder, with $V_v = \pi\ (D/2)^2 \times L_v$.

| Species | $D$ (µm) | $L_V$ (cm) | $T_{PM}$ (nm) | $F_C$ | $F_{PF}$ | $F_P$ | $A_p$ (µm$^2$) | $V_v$ ($10^{-11}$ m$^3$) |
|---|---|---|---|---|---|---|---|---|
| *Corylus avellana* | 19.78±7.17 | 3.55±0.83 | 251±35 | 0.251±0.007 | 0.567±0.026 | 0.142 | 0.314 | $1.09\times10^{-11}$ |
| *Fagus sylvatica* | 16.58±4.96 | 5.15±0.82 | 244±58 | 0.115±0.015 | 0.382±0.106 | 0.044 | 0.118 | $1.11\times10^{-11}$ |
| *Prunus avium* | 16.02±5.08 | 10.6±1.67 | 385±86 | 0.145±0.016 | 0.608±0.023 | 0.088 | 0.470 | $2.14\times10^{-11}$ |

**Table S4** Intercept coefficient (*a*), water potential coefficient (*b*), and temperature coefficient (*c*) for calculating Henry's constant ($K_H$) of different gases (equation 8). Values were obtained through a Multivariate Regression from Table 8 in Mercury *et al.* (2003).

| gas | *a* | *b* | *c* |
|---|---|---|---|
| Ar | 10.0625 | 0.001261 | 0.062230 |
| $N_2$ | 10.8748 | 0.001325 | 0.053611 |
| $O_2$ | 10.1490 | 0.001207 | 0.063000 |
| $CO_2$ | 6.5379 | 0.001302 | 0.107372 |

**Table S5** Values of several variables that were required to model the gas pressure in a recently embolised vessel in flow-centrifuge experiments.

| **Variable** | **Value** | **Units** | **Definition** |
|---|---|---|---|
| $D_p$ | $1.65\times10^{-9}$ | $m^2\ s^{-1}$ | Gas diffusion coefficient in a wet pit membrane |
| $D_w$ | $2\times10^{-9}$ | $m^2\ s^{-1}$ | Gas diffusion coefficient in water |
| $M_{H2O}$ | 18.0153 | $g\ mol^{-1}$ | Molar mass of water |
| $N$ | 500 | - | Number of cylinders used to divide a stem in small compartments, which allowed us to model pressure changes in recently embolised vessels |
| $P_{atm}$ | 101300 | Pa | Atmospheric pressure |
| $R$ | 0.125 | m | Distance from the rotational axis to the downstream end of the centrifuge sample |
| R* | 8.314 | $J\ mol^{-1}\ K^{-1}$ | Gas constant |
| Δt | 0.5 | s | Time interval of the simulation |
| ρ | 997.77 | $kg\ m^{-3}$ | Density of water |

**Table S6** Overview of all abbreviations, units, definitions and equations used in the gas diffusion model for flow-centrifuge measurements.

| Abbreviation | Units | Definition |
|---|---|---|
| $\Delta K_h$ | % | Relative change in hydraulic conductivity (equation 2) |
| $A_p$ | $m^2$ | Total intervessel pit membrane area in a vessel with average dimensions and intervessel connectivity (equation 5) |
| $C_{g\_air}$ | mol $m^{-3}$ | Gas concentration in air (equation 11) |
| $C_{G,i}$ | mol $m^{-3}$ | Partial gas solubility of a gas species "G" in xylem sap in the $i^{th}$ cylinder (equation 9) |
| $C_{h,1}$ | mol $m^{-3}$ | Initial gas concentration in a recently embolised water vapour-filled vessel at time step 1 (equation 13) |
| $C_{h,j}$ | mol $m^{-3}$ | Modelled gas concentration in a recently embolised vessel at time step *j* (equation 20) |
| $C_i$ | mol $m^{-3}$ | Initial gas concentration in the $i^{th}$ cylinder of sap-filled xylem (equation 10) |
| $C_{i,j}$ | mol $m^{-3}$ | Modelled gas concentration in the $i^{th}$ cylinder of sap-filled xylem at time step *j* (equation 16) |
| $C_{relative}$ | % | Relative value of the modelled gas concentration in a recently embolised vessel at time step *j* (equation 21) |
| $D$ | m | Arithmetic mean vessel diameter |
| $F_C$ | % | Intervessel contact fraction, fraction of vessel wall in contact with other vessels as based on transverse sections |
| $F_P$ | % | Pit fraction, fraction of total vessel wall area occupied by intervessel pits |
| $F_{PF}$ | % | Pit-field fraction, fraction of intervessel wall area occupied by intervessel pits |
| $h$ | - | Position of the $h^{th}$ cylinder containing the end wall of a recently embolised vessel at the centre of a centrifuge sample (equation 12) |
| $H^{CC}$ | - | Dimensionless Henry's constant (equation 17) |
| $K_0$ | kg $s^{-1}$ $MPa^{-1}$ $m^{-1}$ | Initial hydraulic conductivity |
| $K_0'$ | kg $s^{-1}$ $MPa^{-1}$ $m^{-1}$ | Hydraulic conductivity before flushing a stem segment |
| $k_a$ | $m^3$ $s^{-1}$ | Axial gas transport rate through xylem sap (equation 14) |
| $K_F'$ | kg $s^{-1}$ $MPa^{-1}$ $m^{-1}$ | Hydraulic conductivity after flushing a stem segment |
| $K_{H,i}$ | bar | Henry's constant for the $i^{th}$ cylinder (equation 8) |
| $K_h$ | kg $s^{-1}$ $MPa^{-1}$ $m^{-1}$ | Hydraulic conductivity |
| $k_p$ | $m^3$ $s^{-1}$ | Axial gas transport rate through the pit membrane (equation 18) |
| $L_V$ | m | Mean vessel length |
| P | Pa | Vapour pressure in a recently embolised vessel |
| PLC | % | Percentage Loss of Conductivity (equation 1) |
| $P_s$ | mol $m^{-3}$ | Partial gas solubility in xylem sap |

| | | |
|---|---|---|
| $r_i$ | m | Distance from the centre of the $i^{th}$ cylinder to the rotational axis |
| T | $^{o}$C | Set temperature in the flow-centrifuge |
| $T_K$ | Kelvin | Set temperature in the flow-centrifuge |
| $T_{PM}$ | m | Intervessel pit membrane thickness |
| $V_v$ | $m^3$ | Volume of one vessel (equation 4) |
| $x$ | m | Distance between the centre of neighbouring cylinders, which also equals the length of the cylinder |
| $\Delta n_{h,j}$ | mol | Amount of gas diffused axially through a pit membrane from the $(h+1)^{th}$ cylinder to a recently embolised vessel at time step $j$ (equation 19) |
| $\Delta n_{i,j}$ | mol | Amount of gas diffused axially through the sap from the $i^{th}$ cylinder to the $(i-1)^{th}$ cylinder at time step $j$ (equation 15) |
| $\rho_{asc}$ | kg $m^{-3}$ | Density of air-saturated water |
| $\Psi$ | MPa | Minimum xylem water potential applied to the middle of a centrifuge sample |
| $\Psi_i$ | MPa | Water potential in the $i^{th}$ cylinder (equation 7) |
| $\omega_{rad}$ | rad $s^{-1}$ | Angular velocity (equation 6) |
| $\omega_{RPM}$ | RPM | The rotational speed of a flow-centrifuge |

**Methods S1** Modelling of the gas pressure in a recently embolised vessel in flow-centrifuge experiments.

To develop a gas diffusion model for flow-centrifuge measurements, we first estimated anatomical traits for branches of the three species studied, and we then divided the model calculation into two parts. We estimated first the initial conditions before an embolism event occurred. Then, we assessed the dynamic changes in the gas concentration between vessels after an embolism event had taken place in the centre of a centrifuge sample.

### Anatomical traits

The three species studied differed in their vessel dimensions (Table S3), which were retrieved from Guan *et al.* (2022). The arithmetic mean vessel diameter (*D*) was measured on semi-thin, transverse sections under a light microscope (Axio Scope A1, Carl Zeiss, Gottingen, Germany).

The pit-field fraction (i.e., fraction of the intervessel wall area occupied by intervessel pits, $F_{PF}$) was based on scanning electron microscopy using tangential surfaces. The intervessel contact fraction (i.e., the fraction of the intervessel wall to the total vessel perimeter, $F_C$) was measured based on transverse sections using a light microscope. The intervessel pit membrane thickness ($T_{PM}$) was measured based on transverse, ultrathin sections observed with a transmission electron microscope (JEM-1400, JEOL Ltd., Tokyo, Japan) at an accelerating voltage of 120 kV. We used a digital camera (F-View, Soft Imaging System GmbH, Münster, Germany) for TEM imaging. The vessel length distribution of the stem was measured with a Pneumatron device (Pereira *et al.*, 2020), and the mean vessel length ($L_V$) was calculated using the equations of Sperry *et al.* (2005). The volume of one vessel ($V_v$) was calculated considering a vessel as a cylinder:

$$V_v = \pi \left(\frac{D}{2}\right)^2 L_V \tag{4}$$

Based on Wheeler *et al.* (2005), the total intervessel pit membrane area in a vessel with average dimensions and intervessel connectivity ($A_p$) was calculated as follows:

$$A_p = \pi \, D \, L_V \, F_C \, F_{PF} \tag{5}$$

**Part 1: Initial gas concentration in the sap**

Here, we assumed that a stem spinning in a flow-centrifuge is symmetric from the edges to the centre of the rotational axis. Thus, the model considered only one side of the stem, from the edge to the centre and, when necessary, we multiplied the calculated values by two. For the simulation, we considered a single vessel with its length and volume depending on the anatomical parameters of the species studied. After an embolism event, we assumed that gas diffused axially through the stem from the edge to the centre of the centrifuge sample, filling this single vessel with gas until it reached atmospheric pressure. To facilitate the calculation, we considered that one side of the stem was composed of one cylindrical vessel, which represents all vessels in a stem with a cross-area $A_p$ and length $R$. Therefore, the gas dissolved in this cylindrical vessel was connected to the gas phase at the edges, which was assumed to be an infinite source of gas. To make the calculations, we divide this cylindrical vessel into $N$ infinitely small cylinders with cross-area $A_p$ and length $x$ (=$R/N$). Considering that these small cylinders had a length close to zero, we assumed that the conditions along the $i^{th}$ cylinder were homogenous.

Then, we calculated the initial condition along the spinning sample, before the embolism took place. Taking into account that the pressure increases along the axis towards the outer sides of the sample (Alder *et al.*, 1997), the initial gas concentration in the $i^{th}$ cylinder of xylem sap ($C_i$) was affected by both the rotational speed of the flow-centrifuge and the distance from its centre to the rotational axis ($r_i$). Therefore, based on Alder *et al.* (1997) and the flow-centrifuge rotational speeds ($\omega_{RPM}$) applied in experiment 2 (Table S1), the water potential in the $i^{th}$ cylinder ($\Psi_i$) was calculated as follows:

$$\omega_{rad} = \frac{2\pi\ \omega_{\mathrm{RPM}}}{60} \tag{6}$$

$$\Psi_i = -\frac{0.5\rho\omega_{rad}^2(R^2 - {r_i}^2)}{10^6} \tag{7}$$

Where ρ was the density of water in g mol$^{-1}$; $\omega_{rad}$ was the angular velocity of the flow-centrifuge in rad s$^{-1}$, and $R$ was the distance (in m) from the rotational axis to the distal (downstream) end of the centrifuge sample.

Considering the water potential ($\Psi_i$) calculated by equation 7 and the temperature (T) applied in experiment 2, Henry's constant for the $i^{th}$ cylinder ($K_{H,i}$, in bars) was estimated according to Mercury *et al.* (2003) and Schenk *et al.* (2016) as follows:

$$K_{H,i} = e^{\left(\frac{a+b\psi_i+c\mathrm{T}}{\ln(\mathrm{T})}\right)} \tag{8}$$

Where *a*, *b* and *c* were the intercept coefficient, the water potential coefficient, and the temperature coefficient, respectively. These values were obtained through a Multivariate Regression from table 8 in Mercury *et al.* (2003), and shown in Table S4.

Considering $\Psi_i$ and T, the density of water (ρ) was corrected for the density of air-saturated water ($\rho_{asc}$) according to Jones & Harris (1992). Then, based on Sander (2015), the partial gas solubility of a gas species "G" in xylem sap in the $i^{th}$ cylinder ($C_{G,i}$) was calculated as follows:

$$C_{G,i} = 10\frac{P_s \rho_{asc}}{K_{H,i} M_{H_2O}} \tag{9}$$

Where $P_s$ was the partial gas solubility, and $M_{H_2O}$ was the molar mass of water.

We considered that the atmosphere was composed of Ar, $N_2$, $O_2$ and $CO_2$ (0.93, 78.09, 20.98, 0.10% respectively), and estimated $C_i$ as the sum of $C_{G,i}$ as follows:

$$C_i = C_{Ar,i} + C_{N_2,i} + C_{O_2,i} + C_{CO_2,i} \tag{10}$$

The gas concentration in air ($C_{g\_air}$) at atmospheric pressure ($P_{atm}$ = 101300 Pa) was calculated as:

$$C_{g\_air} = \frac{P_{atm}}{R^* \, T_K} \tag{11}$$

Where $R^*$ was the gas constant (8.314 J $mol^{-1}$ $K^{-1}$) and $T_K$ was the air temperature in Kelvin.

**Part 2: Dynamic changes in the gas concentration in a recently embolised vessel**

Using the anatomical traits of the species studied and the initial gas concentration in the sap, we simulated an embolised water vapour-filled vessel at the centre of the centrifuge sample at time zero (t=0) and assessed the dynamic changes in gas concentration. Once the equilibrium between the gas and liquid phase in xylem sap was broken by a recently embolised vessel, gas diffusion started to gradually fill this embolised vessel with gas (Wang *et al.*, 2015). Here, we only considered axial gas diffusion, which was estimated over time as described by equations 9 and 10 in Yang *et al.* (2023). We assumed that radial diffusion was much slower (Wang *et al.*, 2015) and would not contribute to the gradual gas increase in the embolised vessel considering the 1 hour duration of the experiments. By using gas diffusion rates, we also estimated the changes in the gas concentration of the *N* cylinders of xylem as well as the recently embolised vessel. The simulation calculated the gas concentrations every 0.5 seconds ($\Delta t$) for the duration of 1 hour. Thus, the simulation was made in 7200 steps over 1 hour.

Herein, we assumed that the embolised vessel was a cylinder with a volume $V_V$. In this case, the entire vessel was located at the centre of the rotational axis ($i$ = 1). This means that half of the vessel extended from the centre to the position of the $h^{th}$ cylinder as follows:

$$h = \frac{\frac{L_V}{2}}{x} \quad (12)$$

Where $x$ was the length of a cylinder, and $L_V$ was the mean vessel length.

The recently embolised vessel was initially filled with water vapour (P=3200 Pa according to Wang *et al.* [2015]). We assumed that the conditions in the entire embolised vessels were homogenous. Thus, the first *h* cylinders covered half of the vessel, and had the same gas concentration. The initial gas concentration in a recently embolised vapour-filled vessel at time step 1 ($C_{h,1}$) was calculated as follows:

$$C_{h,1} = \frac{P}{R^{*}\ T_K} \quad (13)$$

Where $R^{*}$ was the gas constant (8.314 J $mol^{-1}$ $K^{-1}$) and $T_K$ was the air temperature in Kelvin.

We assumed that gas diffusion in the centre of a centrifuge sample (i.e., between the $(h+1)^{th}$ and $h^{th}$ cylinder) occurred through a pit membrane. We then calculated the gas diffusion through the sap between the *N*-$h$ cylinders. We considered that the gas diffusion coefficient in water ($D_w$) was $2\times10^{-9}$ m$^2$ s$^{-1}$, and calculated the axial gas transport rate through the sap ($k_a$, in m$^3$ s$^{-1}$) as follows:

$$\mathrm{k}_a = \frac{A_\mathrm{p}\mathrm{D}_\mathrm{w}}{x} \tag{14}$$

Where $A_p$ was the total intervessel pit membrane area, and $x$ was the distance between the centre of neighbouring cylinders and was also the length of the cylinder.

The amount of gas diffused axially through the sap from the $i^{th}$ cylinders to the $(i\text{-}1)^{th}$ cylinders at time step $j$ ($\Delta n_{i,j}$, in mol) was calculated as follows:

$$\Delta\mathrm{n}_{i,j} = \mathrm{k}_\mathrm{a}(\mathrm{C}_{i,j} - \mathrm{C}_{i-1,j})\Delta\mathrm{t} \tag{15}$$

Where Δt was the time interval of gas diffusion.

The modelled gas concentration in the $i^{th}$ cylinder of xylem sap at the time step $j$ ($C_{i,j}$) was then calculated as:

$$\mathrm{C}_{i,j} = \mathrm{C}_{i,j-1} + \frac{\Delta\mathrm{n}_{i,j} - \Delta\mathrm{n}_{i+1,j}}{A_p x} \tag{16}$$

Where $x$ was the length of the cylinder.

Considering the initial gas concentration in the $h^{th}$ cylinder of xylem sap ($C_h$) calculated by equation 10, the dimensionless Henry's constant ($H^{CC}$) was estimated as follows:

$$\mathrm{H^{CC}} = \frac{\mathrm{C}_h}{\mathrm{C}_{g\_air}} \tag{17}$$

Where $C_{g\_air}$ was the gas concentration in air at atmospheric pressure.

Here, we considered that the gas diffusion coefficient across a wet pit membrane ($D_p$) was $1.65\times10^{-9}$ $m^2$ $s^{-1}$, and calculated the axial gas transport rate through an intervessel pit membrane ($k_p$, in $m^3$ $s^{-1}$) as follows:

$$\mathrm{k_p} = \frac{A_p \mathrm{D_p} H^{CC}}{T_{PM}} \tag{18}$$

Where $T_{PM}$ was the intervessel pit membrane thickness in m.

The amount of gas diffused axially through the pit membrane from the cylinder $(h+1)^{th}$ to the recently embolised vessel at the time step $j$ ($\Delta n_{h,j}$, in mol) was calculated as follows:

$$\Delta \mathrm{n}_{h,j} = \mathrm{k_p}(\mathrm{C}_{h+1,j} - \mathrm{C}_{h,j}\mathrm{H}^{CC})\Delta t \tag{19}$$

The modelled gas concentration in the recently embolised vessel at the time step $j$ ($C_{h,j}$) was then calculated as:

$$\mathrm{C}_{h,j} = \mathrm{C}_{h,j-1} + \frac{2\Delta n_{h,j}}{V_v} \tag{20}$$

**Modelled gas concentration in a recently embolised vessel versus hydraulic conductivity measurements**

Using the model variable values described in Table S5 and considering the anatomical traits (Table S3), the temperature and the water potentials applied to flow-centrifuge experiments (Table S1), we calculated $C_{h,j}$. However, the absolute values of $C_{h,j}$ varied according to temperature. Thus, to facilitate comparison between species and different water potentials as well as to correlate $C_{h,j}$ to measured hydraulic conductivity, we considered the relative values of $C_{h,j}$ ($C_{relative}$), taking as reference the maximum and minimum of $C_{h,j}$. We assumed that $C_{h,1}$, and $C_{g\_atm}$ represented the minimum and maximum value of $C_{h,j}$, respectively, and calculated $C_{relative}$ as follows:

$$C_{relative} = 100\frac{(C_{h,j} - C_h)}{(C_{g\_atm} - C_h)} \quad (21)$$

All analyses and models were developed using the R programming environment. Codes are available upon request.

**Methods S2** Multiple regression for $\Delta K_h$ as a function of time and $\Psi$.

In our trial measurements, we noticed that the curve $\Delta K_h$ versus time showed a logarithmic pattern for each $\Psi$. We also noticed that the scale of $\Delta K_h$ changed according to $\Psi$. Thus, there was an interdependence between $\Delta K_h$ and time and $\Psi$. Hence, $\Psi$ affected the coefficient of the equation, which described the relationship between $\Delta K_h$ and time. Taking these findings into account, the multiple non-linear regression was made in two steps. First, we made a regression between $\Delta K_h$ and time for each $\Psi$. We transformed time (t) using the natural logarithm (ln) and found the slope ($\alpha$) and intercept coefficient ($\beta$) for each $\Psi$, as follows:

$$\Delta K_h(\Psi, t) = \alpha(\Psi)\ln(t) + \beta(\Psi) \quad (22)$$

In the second step, we made non-linear and linear regressions to find the equations that describe α and β as a function of Ψ. For the non-linear regression, we considered the smallest n-degree equation, which provided an $R^2$ higher than 0.65.

$$\alpha(\Psi) = a_i\Psi^n + a_{i-1}\Psi^{n-1} + a_{i-2}\Psi^{n-2} + \ldots a_1\Psi + a_0 \tag{23}$$

$$\beta(\Psi) = \mathrm{b}_1\Psi + b_0 \tag{24}$$

Once we calculated $a_n$, $a_{n-1}$,…,$a_0$, and the $b_1$ and $b_0$ coefficients, we found the equation that described $\Delta K_h$ as a function of Ψ and t as follows:

$$\Delta\mathrm{K_h}(\Psi, \mathrm{t}) = (a_i\Psi^n + a_{i-1}\Psi^{n-1} + a_{i-2}\Psi^{n-2} + \ldots a_1\Psi + a_0\ln(\mathrm{t}) + (\mathrm{b}_1\Psi + b_0) \tag{25}$$

All these regressions were performed once for each species at 22°C. The analyses were developed using Excel and are available upon request.

**Methods S3** Fitting a rectangular hyperbola model.

Based on the non-linear regression models of Archontoulis & Miguez (2015), we selected the rectangular hyperbola model as the most suitable model and fitted it for the relationship between hydraulic conductivity ($K_h$) and time. The expression for a rectangular hyperbola takes into account both the maximum height of the asymptote and the rate of approach towards it, and was written as follows:

$$\mathrm{y(x)} = \frac{\mathrm{ax}}{\mathrm{b + x}} \tag{26}$$

Where y was the dependent variable, x was the independent variable, a was the asymptote, and b was the parameter defining the rate of approaching the asymptote.

Considering x = t and making the following transformation, we were able to fit the model and found the equation that described $K_h$ as a function of time:

$$y(t) = -K_h(t) + K_0 \tag{27}$$

$$K_h(t) = -\frac{at}{b + t} + K_0 \tag{28}$$

Where $K_0$ is the hydraulic conductivity measured at time zero.

Therefore, the minimum value of hydraulic conductivity ($K_{min}$) predicted by the rectangular hyperbola model could be estimated as follow:

$$K_{min} = K_0 - a \tag{29}$$

All analyses and models were developed using the R programming environment, specifically the package ‘drc’. Codes are available upon request.

**Box S1** Workflow of the axial gas diffusion model for flow-centrifuge experiments. The model was based on the Unit Pipe Pneumatic model of Yang *et al.* (2023) and estimated the axial gas diffusion through the sap and pit membranes after an embolism event in the centre of the centrifuge sample. This model was developed to evaluate the functional link between the gas concentration in a recently embolised, vapour-filled vessel and hydraulic conductivity in flow-centrifuge measurements. The abbreviations are described in Methods S1, and summarized in Table S6.

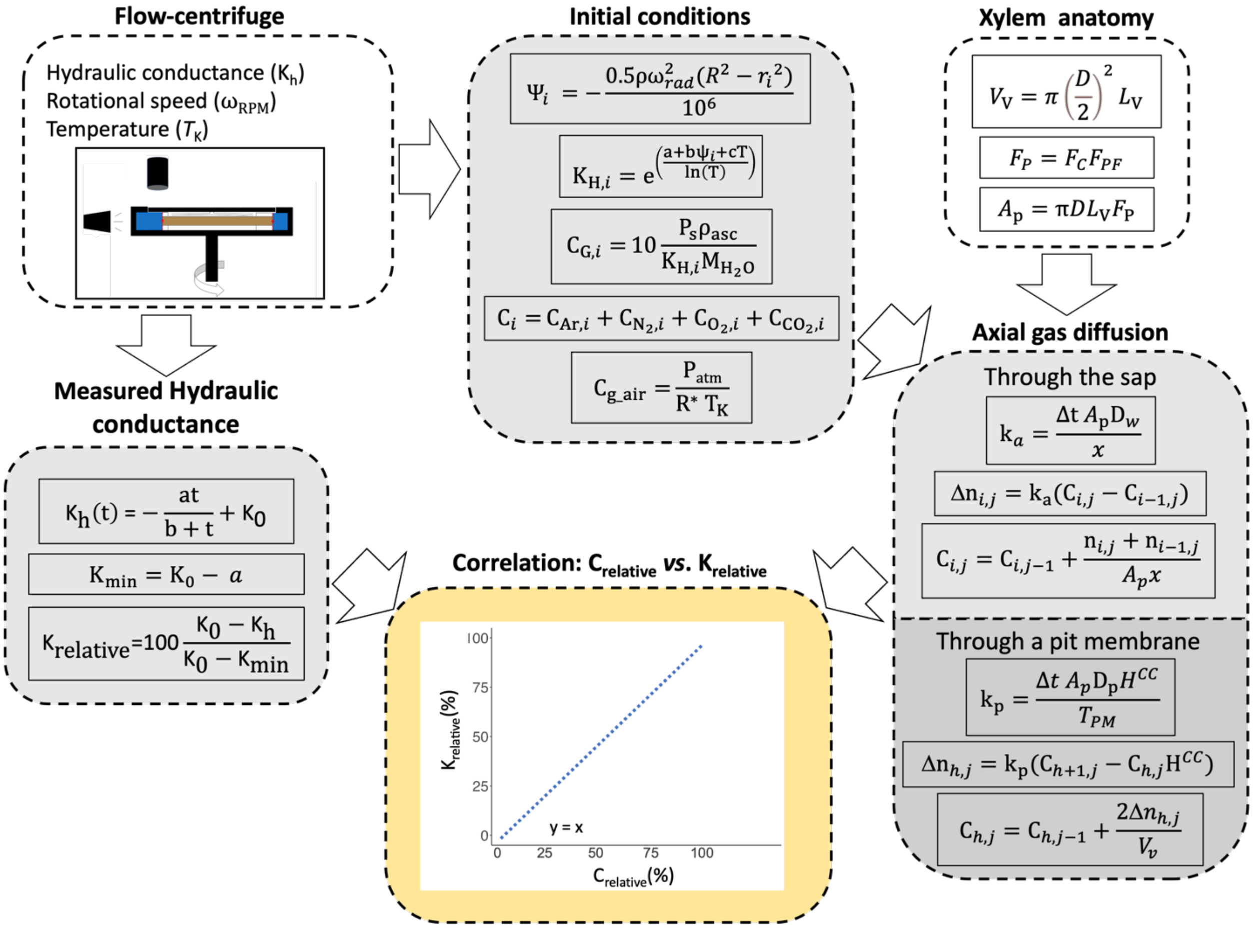
